# Luminal Vascular Dysfunction Drives Rapid Blood Brain Barrier Injury in Hyperglycemic Stroke: Key Roles for Luminal Glycocalyx and Complement

**DOI:** 10.1101/2025.06.26.661858

**Authors:** Hansen Chen, Jacqueline A. Frank, Chunfeng Tan, Alex G. Lee, Richard Kopchock, Terrance Chiang, Anika Kim, Manuel Galvan, Justin F. Fraser, David Dornbos, Hassan Aboul-Nour, Nathan Millson, Stephen Tomlinson, Louise D. McCullough, Keith Pennypacker, Michelle Y. Cheng, Tonya M. Bliss, Gary K. Steinberg

## Abstract

**Background:** Acute hyperglycemia affects approximately 40% of stroke patients and is associated with worse outcomes. The underlying mechanisms linking this metabolic stress to stroke-induced brain injury remains unclear, and effective therapies are lacking.

**Methods:** In a mouse model of acute hyperglycemic stroke, luminal disruption, blood-brain barrier (BBB) leakage, neurological deficit, motor function, and mortality were evaluated. Vascular luminal glycocalyx and complement activation were assessed by immunostaining, with glycocalyx loss confirmed by electron microscopy. Complement C3’s causal role was tested using C3 knockout mice and site-targeted inhibition with CR2-Crry. To enhance translational relevance, post-mortem human stroke and control brains were immunostained to assess the association between endothelial glycocalyx loss and vascular complement activation. In a separate stroke patient cohort, soluble complement activation products were measured in pre-thrombectomy plasma, and their predictive value for modified Rankin Scale (mRS) outcomes evaluated using elastic net regression.

**Results:** Hyperglycemic stroke mice exhibited accelerated and more severe BBB breakdown, greater functional deficits, and higher mortality than normoglycemic controls, mirroring clinical observations. Acute hyperglycemia triggered rapid vascular luminal injury characterized by loss of endothelial luminal glycocalyx, luminal IgM/IgG deposition, and vascular complement C3 activation, leading to BBB disruption. This vascular luminal injury was corroborated in human stroke brain tissue. These luminal changes persisted despite glucose normalization and were exacerbated by reperfusion, driving injury into the brain parenchyma. Genetic and pharmacological approaches confirmed vascular complement activation as a causal driver of severe BBB disruption and poor outcomes. Importantly, site-targeted pharmacological inhibition of complement <u>after</u> reperfusion preserved BBB integrity and improved outcomes, defining a time-specific, luminal-directed strategy as a promising adjunct to thrombectomy. Notably, soluble complement activation markers in pre-thrombectomy stroke plasma predicted clinical outcomes, highlighting their potential as pre-intervention markers for patient stratification and tailored therapy.

**Conclusion:** This study reframes acute hyperglycemic stroke as a vascular luminal disorder, establishing a novel **Metabolic–Complement–Vascular (MCV) axis** linking metabolic stress to endothelial luminal glycocalyx loss, vascular complement activation, and BBB breakdown in both mice and humans. This new mechanistic understanding transforms the therapeutic landscape of hyperglycemic stroke, offering a potential time-defined, luminal-focused adjunct therapy alongside thrombectomy.

**Clinical Perspective:** *What Is New?:*

- This study reframes hyperglycemic stroke as an acute vascular **luminal** problem, marked by rapid loss of endothelial luminal glycocalyx and complement C3 activation at the vascular luminal surface.
- The rapid luminal changes, identified in both rodent and human stroke brain tissues, establish a novel **Metabolic–Complement–Vascular (MCV) axis** linking metabolic stress to luminal damage, blood-brain barrier (BBB) breakdown, and injury progression into the brain parenchyma.
- The first clinical evidence that pre-thrombectomy plasma complement activation markers independently predict stroke outcomes — laying the foundation for risk stratification before reperfusion and precision adjunct therapies.

*What Are the Clinical Implications?:*

- The rapidity and persistence of the MCV axis activation—even after glucose normalization—help explain the limited efficacy of insulin therapy, shifting the therapeutic focus from glycemic control to luminal-targeted interventions.
- Identification of a narrow but actionable window following reperfusion where complement C3 inhibition preserves the BBB and limits injury progression to the parenchyma, offers a promising adjunct to thrombectomy in metabolically vulnerable stroke patients.
- Targeting the vascular luminal surface reshapes the therapeutic landscape for hyperglycemic stroke by enabling systemic interventions—bypassing the challenge of BBB penetration, supporting rapid clinical translation using existing FDA-approved C3 inhibitors, and framing the endothelial glycocalyx as a promising area for therapeutic and diagnostic exploration in hyperglycemic stroke and broader cerebrovascular disease.

## Introduction

Ischemic stroke, caused by blockage of a blood vessel in the brain, is a leading cause of death and disability worldwide^1,2^. Although reperfusion therapies to restore blood flow, such as thrombectomy and thrombolysis, have transformed acute stroke care, fewer than half of treated patients achieve favorable outcomes^3^. Acute hyperglycemia, a transient spike in blood glucose that occurs in approximately 40% of ischemic stroke patients, irrespective of whether they have diabetes^4^, is a key predictor of poor outcomes following these interventions^5–7^. Attempts to manage this risk with insulin administration have not yielded consistent clinical benefit^8^, and no standard adjunctive treatment currently exists for this high-risk group—highlighting a critical unmet medical need. This emphasizes the importance of uncovering the underlying mechanisms of hyperglycemic injury to identify targeted interventions.

Both clinical and preclinical studies consistently show that acute hyperglycemia worsens stroke outcomes and exacerbates blood–brain barrier (BBB) disruption^9^. While rodent studies have provided valuable mechanistic insights including oxidative stress^10–12^, matrix metalloproteinase (MMP) activation^13^, and detrimental macrophage polarization^14^, they have primarily focused on processes within the brain parenchyma and at the later phases of injury, starting at 24 hours after stroke. However, the limited success of glucose correction in clinical settings within this same timeframe suggests these brain parenchymal mechanisms are not the primary drivers of hyperglycemic stroke injury, and that earlier mechanisms are more important. This prompted a shift in perspective: we propose that since a transient surge in glucose is a systemic stressor, it targets the luminal side of the brain vasculature, rapidly initiating a cascade of pathological events. The specific vascular events linking peripheral hyperglycemia to exacerbated stroke brain injury, however, remain unknown.

The luminal surface of the brain vasculature is coated by the endothelial glycocalyx, a carbohydrate-rich layer of glycoproteins and glycans that forms the first line of defense between the blood stream and brain parenchyma^15^. This specialized structure functions as a physical barrier and an immune sensor, playing a critical role in maintaining vascular integrity and BBB function^15,16^. While acute hyperglycemia has been shown to alter the glycocalyx in the peripheral vasculature^17^, the glycocalyx in the brain is thicker, has a different glycoprotein composition^18^, and appears to be more resilient against stressors such as lipopolysaccharide (LPS)^19^. However, the impact of acute hyperglycemia, as a metabolic stressor, on this specialized luminal interface during stroke remains unknown. Understanding whether glycocalyx dysfunction contributes to the rapid and severe BBB damage observed in hyperglycemic stroke could be essential for preventing metabolic brain injury.

While glycocalyx loss compromises vascular integrity, it may also act as a catalyst for immune activation. Prior studies in peripheral vessels suggest that glycocalyx loss coincides with complement activation within the vessels^20^. However, whether a similar link exists at the blood–brain barrier remains unknown. While complement has been studied in stroke, particularly for its role in modulating glia cells, infarct growth, and neurological injury under normoglycemic conditions ^21–23^, most studies have focused on the brain parenchyma, with little attention on vascular complement, especially in the setting of systemic metabolic stress. It remains unclear whether complement C3 activation directly contributes to BBB disruption, or whether its activation originates in the brain or at the vascular interface—a distinction that may differ fundamentally between normoglycemic and hyperglycemic stroke. Notably, the hyperacute phase, a likely key window in hyperglycemic injury, has been largely overlooked. Addressing these gaps is essential to identify drivers of hyperglycemia-induced vascular injury and to advance complement-targeted therapeutics.

Here, we investigate how acute hyperglycemia disrupts the brain’s vascular interface, activates complement, and contributes to BBB injury after stroke. We propose the concept of a metabolic–complement–vascular (MCV) axis linking systemic metabolic stress to cerebrovascular injury, particularly the luminal glycocalyx, and highlight a critical window for intervention. Using experimental stroke models and human brain and plasma analyses, we map the temporal and spatial dynamics of this process and identify the vessel lumen, particularly vascular complement activation, as a potential adjunct therapeutic target, in conjunction with thrombectomy, to improve outcomes in hyperglycemic stroke.

## Results

### Hyperglycemic Stress Worsens Stroke Outcomes and Induces Severe, Pathologic BBB Disruption

Acute hyperglycemic stroke was induced by systemic injection of glucose 10 minutes prior to transient middle cerebral artery occlusion (MCAO) (Figure 1A), according to an established protocol^24^. Glucose administration caused an approximately twofold increase in blood glucose compared to baseline (Figure 1B) — a magnitude of elevation comparable to stress hyperglycemia observed in stroke patients, particularly in nondiabetics^25^. Blood glucose levels returned to baseline within two hours (Figure 1B). This transient hyperglycemia (HG) significantly worsened stroke outcomes, mirroring clinical observations. All hyperglycemic mice exhibited progressive weight loss and died within 8 days post-stroke (Figure 1C, D). In contrast, normoglycemic mice began to gain weight by 4 days post-stroke and 75% survived to the end of the study (day 14) (Figure 1C, D). Neurological deficit scoring revealed significantly worsened functional outcomes in hyperglycemic mice on days 1, 3, and 7 after stroke (Figure 1E), further supported by worse general deficit scores (Figure S1A). These findings indicate that even a short episode of acute hyperglycemia at the time of stroke can have lasting detrimental effects on survival and neurological function.

**Figure 1.**
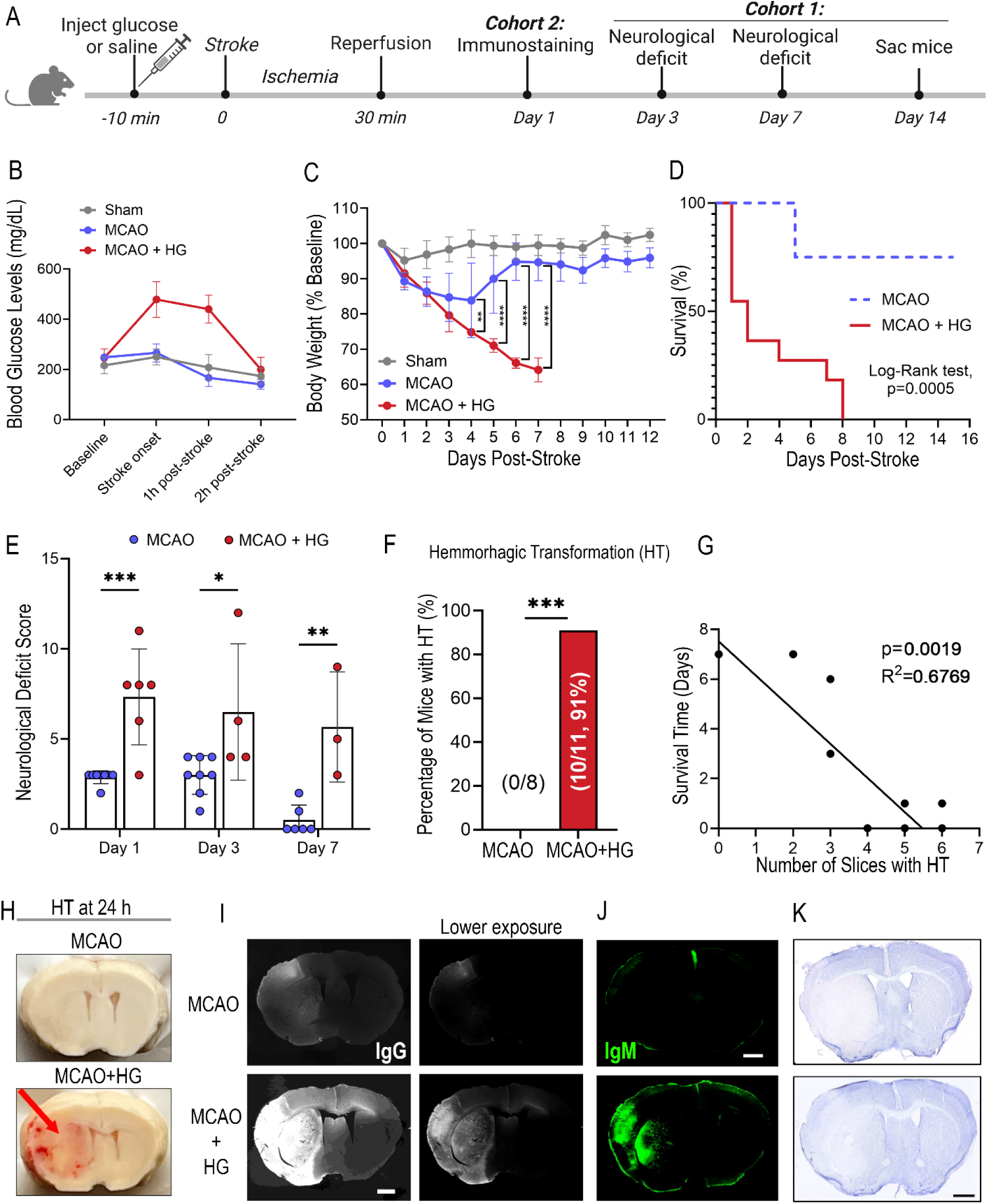
Acute hyperglycemia worsens stroke outcome and causes severe blood-brain barrier disruption. **A**, Schematic of the experimental timeline. Mice received intraperitoneal glucose (hyperglycemia; HG) or saline 10 minutes prior to transient middle cerebral artery occlusion (MCAO), followed by reperfusion, and were monitored over 14 days (cohort 1) or sacrificed at 24-hour post-stroke for histological analysis. Cohort1: n=8-11/group. Cohort 2: n=5/group. **B**, Blood glucose levels measured at indicated time points relative to stroke. Glucose levels were significantly elevated in the MCAO + HG group at early time points and returned to normal within 2 hours. n=5/group. **C**, Body weight trajectories post-stroke. **p<0.01; ****p<0.0001. **D**, Kaplan–Meier survival curve showing acute hyperglycemia significantly reduces survival after stroke (Log-rank test, p = 0.0005). **E**, Neurological deficit scores on post-stroke day 1, 3, and 7. ***p<0.001; ****p<0.0001. **F**, Percentage of mice exhibiting hemorrhagic transformation (HT) at post-mortem analysis during the 14 days post-stroke. ***p<0.001, Fisher’s exact test. **G**, Survival time of individual hyperglycemic mice negatively correlated (Pearson) with the total number of hemorrhagic brain slices identified by post-mortem brain analysis. **H**, Representative brain images showing hemorrhagic transformation (HT) at 24 hours post-stroke (cohort 2). n=5-7. **I**, Representative images of brain IgG immunostaining at 24 hours post-stroke, shown at both normal and reduced exposure levels. n=5. Scale bar: 1000µm. **J**, Representative images of brain IgM immunostaining at 24 hours post-stroke. n=5. Scale bar: 1000µm. **K**, Representative Cresyl Violet staining of brain sections at 24 hours post-stroke, showing no difference in infarct size between the normoglycemia and hyperglycemia groups. n=5-6. Scale bar: 1000µm. Data are shown as mean ± SD.

Clinically, hyperglycemia is associated with increased risk of hemorrhagic transformation (HT) after stroke, a hallmark of severe vascular injury. Consistent with this, post-mortem analysis (Figure 1; Cohort 1) of brains revealed severe HT in hyperglycemic mice, with 91% exhibiting macroscopic HT, whereas normoglycemic mice did not develop HT (Figure 1F; Figure S1B). Notably, the total number of hemorrhagic brain slices inversely correlated with days of survival(R² = 0.6769, p = 0.0019; Figure 1G), suggesting that HT burden and severe vascular injury contribute to early mortality. Successful infarction was confirmed in all mice included, either by 2,3,5- Triphenyltetrazolium chloride (TTC) for mice that died before the study endpoint, or by CD68 and GFAP staining (Figure S1 B, C).

Mice in the first cohort died at different timepoints, therefore, to characterize early-stage effects of hyperglycemia more precisely, we conducted a separate experiment (Cohort 2) in which all mice were sacrificed at 24 hours post-stroke. Hyperglycemic mice showed highly exacerbated vascular permeability compared to normoglycemic mice as evidenced by pronounced HT (Figure 1H), increased hemispheric swelling (Figure S1E), and markedly elevated IgG (a known marker of BBB leakage) staining (Figures 1I, Figure S1F), all indicative of significantly greater BBB leakage. The IgG levels in hyperglycemic mice were so elevated that imaging at a standard exposure, appropriate for detecting IgG leakage in normoglycemic mice, resulted in a saturated signal in hyperglycemic mice; therefore, all further imaging was conducted at a lower exposure tailored to the hyperglycemic mice (Figure 1I).

Notably, the immune protein IgM, which is about six times larger than IgG (∼900 kDa vs. 150 kDa), showed marked extravasation in hyperglycemic stroke mice but was almost undetectable in normoglycemic stroke mice brains (Figure 1J, Figure S1G). This further highlights the extensive disruption of the BBB due to acute hyperglycemia. Infarct size, by contrast, was not significantly altered by hyperglycemia (Figure 1K, Figure S1D), suggesting that overwhelming BBB disruption, rather than infarct expansion, is the primary mechanism through which hyperglycemia exacerbates stroke outcome.

### Rapid and Severe Disruption of Vessel Lumen Integrity in Hyperglycemic Stroke

Given the transient nature of acute hyperglycemia, we next examined whether vascular injury begins even earlier at the luminal surface—the frontline directly exposed to circulating glucose. We focused on the 4.5-hour post-stroke window (Figure 2A), a critical period for acute intervention.

**Figure 2.**
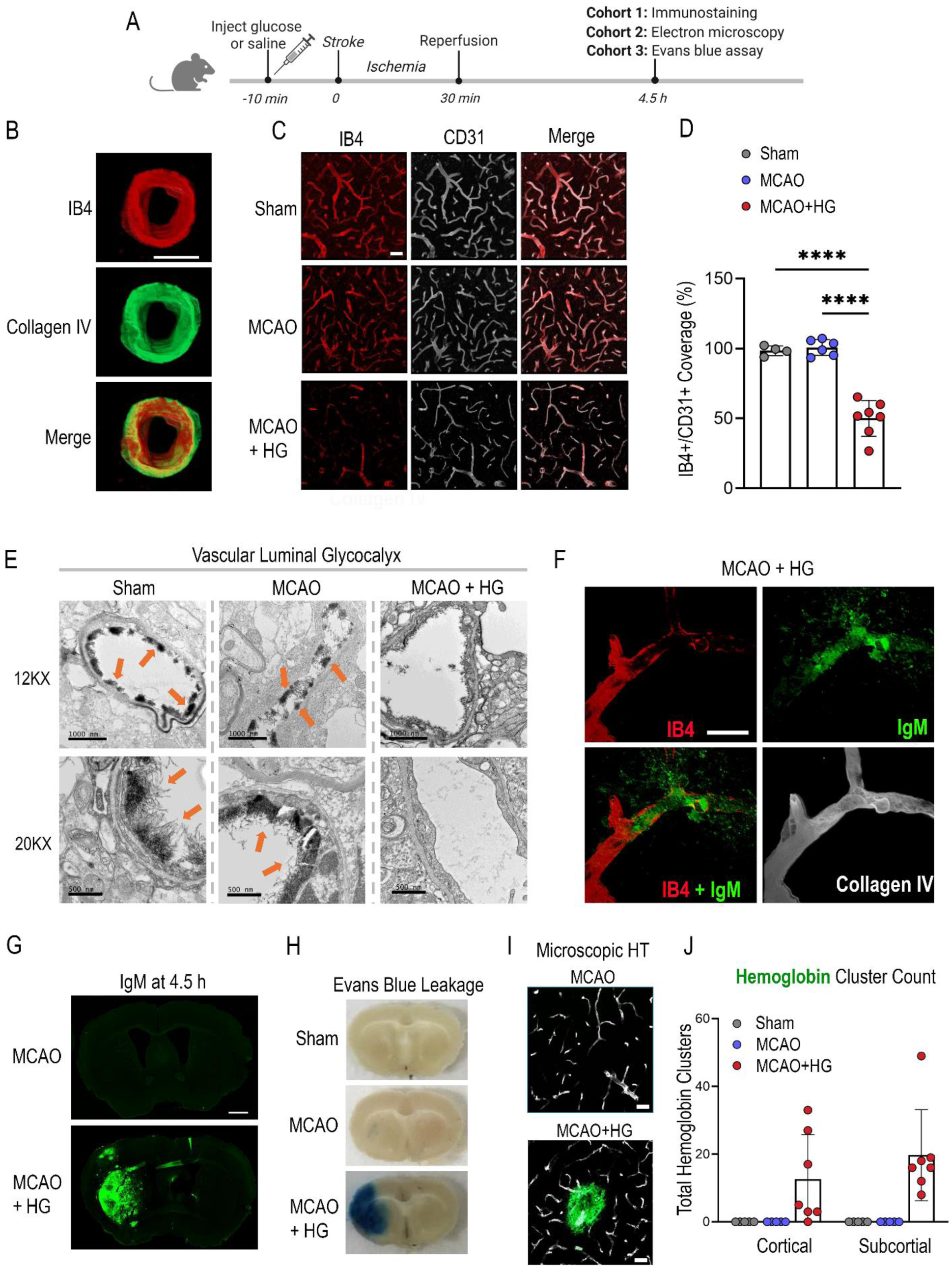
Acute hyperglycemia rapidly disrupts endothelial luminal integrity and induces early blood-brain barrier damage. **A**, Schematic of the experimental timeline. **B**, Co-staining of IB4 and collagen IV showing that IB4 predominantly labels the luminal side of brain vessels. Scale bar: 10µm. **C**, Immunostaining of brains at 4.5h post-stroke with IB4 lectin (red), to label luminal glycoproteins, and CD31 endothelial marker (white). n=5-7/group. Scale bar: 40µm. **D**, Ratio of the area covered by IB4 staining to the area covered by CD31 staining in 3 images per brain. n=5-7/group. **E**, Representative transmission electron microscopy images of cortical vessels in the ischemic territory. Arrows indicate luminal glycocalyx structure. **F**, Immunofluorescent staining showed IgM accumulation around disrupted vessels, co-localizing with loss of IB4 signals in CD31+ brain vessels in MCAO + HG mice, at 4.5 hours post-stroke. n=5. Scale bar: 20µm. **G**, Representative IgM immunostaining in MCAO and MCAO + HG mice brains at 4.5 hours post-stroke. n=5/group. Scale bar: 1000µm. **H**, Representative brain images showing significantly increased Evans blue leakage in the ipsilateral hemisphere of hyperglycemic mice (MCAO + HG). n=5/group. **I-J,** Hemoglobin staining (green, **I**) and quantification (**J**) revealed early microscopic hemorrhagic transformation (HT) in the stroke territory of MCAO + HG mice. n=5-7/group. Scale bar:40 µm. Data are presented as mean ± SD. One-way ANOVA with Tukey’s post hoc test was used for multiple comparisons. ****p < 0.0001.

Staining for IB4 lectin, which preferentially binds to luminal glycoprotein-rich structures (Figure 2B, Figure S2C)^26^ revealed a significant loss of IB4 signal in MCAO + HG mice (versus normoglycemic stroke: Figure 2C-D), suggesting rapid and substantial luminal disruption. This finding was further substantiated by ultrastructural analysis of the brain vessels via transmission electron microscopy, which demonstrated a rapid loss of the endothelial luminal glycocalyx in hyperglycemic mice, with near-complete loss of this protective layer; this is in stark contrast to the preserved glycocalyx observed in normoglycemic controls at this early 4.5-hour time point (Figure 2E). Notably, vessel segments with reduced IB4 staining (i.e. disrupted luminal integrity) exhibited greater IgM accumulation within the lumen, along with increased perivascular IgM deposition indicative of vessel leakage (Figure 2F).

This luminal vascular vulnerability translates into striking differences in early BBB disruption between the normoglycemic and hyperglycemic stroke. Hyperglycemic mice exhibited marked IgM leakage into the brain parenchyma, whereas normoglycemic controls showed minimal to no signal—indicating that BBB disruption under hyperglycemic stress is already severe enough at this early stage to permit passage of large immune proteins such as IgM (Figure 2G). To further support this, Evans Blue dye extravasation assays showed markedly increased Evans Blue accumulation in the ipsilateral hemisphere of hyperglycemic stroke mice, compared to minimal leakage in normoglycemic stroke (Figures 2H, Figure S2A). Evans Blue levels in the liver were comparable across groups, confirming consistent tracer delivery (Figure S2B). Additionally, microscopic HT, identified by prominent perivascular hemoglobin clusters, was observed exclusively in the hyperglycemic group even at this early time point (Figures 2I–J), further confirming rapid and severe BBB disruption under hyperglycemic stress. Together, these findings show that even a brief episode of acute hyperglycemia can rapidly and severely compromise vascular luminal architecture, driving immune protein deposition and barrier disruption within hours of stroke onset, emphasizing the importance of targeting upstream drivers of BBB injury in this early time window.

### Rapid Vascular Complement Activation Precedes BBB Breakdown in Hyperglycemic Stroke

Since IgM, along with IgG, is a potent activator of the complement pathway^27^, we investigated whether the presence of IgM at the disrupted luminal surface (Figure 2F) is associated with early complement activation under hyperglycemic stress. To evaluate this, we stained for C3d, a surface-bound activation product of the key complement protein C3, that remains at the site of activation. As severe BBB leakage was already evident by 4.5 hours post-stroke (Figure 2), we conducted a time-course analysis at earlier stages to capture upstream molecular events. This revealed that hyperglycemic stroke mice exhibited very rapid and robust vascular complement activation in the ischemic territory, with obvious C3d deposition detectable at 1 hour after stroke onset, progressively increasing over time (Figure 3A–B). In contrast, normoglycemic stroke mice exhibited negligible C3d expression at any time point (Figure S3).

**Figure 3.**
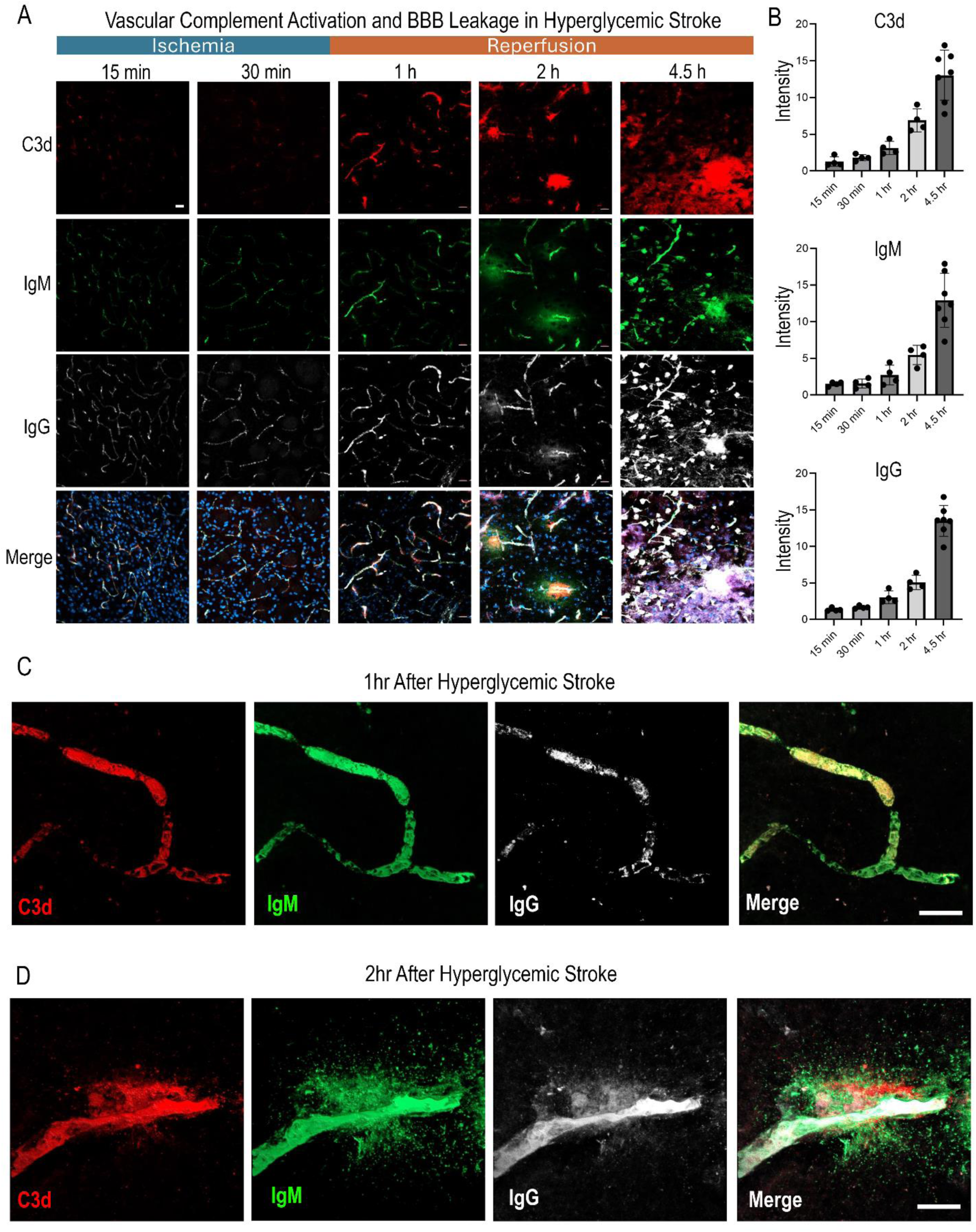
Time course showing rapid complement activation and immunoglobulin deposition/leakage after hyperglycemic stroke. **A**, Post-stroke time-course immunostaining of C3d, IgM, and IgG in brain sections from MCAO + HG mice. n=4- 7/group. Scale bar: 20µm. **B**, Quantification of C3d, IgM, and IgG signal intensity across time points (n = 4-7/group). Data: mean ± SD. **C**, 3D confocal images showing vascular co-localization of IgG, IgM, and C3d in brain vessels in MCAO + HG mice at 1 h post- stroke. Scale bar: 20 µm. **D**, IgG, IgM, and C3d deposition extended beyond blood vessels into adjacent brain parenchyma at 2 h post-stroke. Scale bar: 20 µm.

Two distinct phases of complement activation were identified. The initial phase occurred very rapidly within the vasculature, with IgG, and IgM detected in the vasculature as early as 15 minutes after stroke onset, accompanied by faint, punctate C3d deposition, suggesting the onset of vascular complement activation (Figure 3A-B). By 1-hour post-stroke (30 minutes after reperfusion), all signals intensified, indicating an escalating intravascular complement response. High-resolution confocal imaging at 1-hour post-stroke confirmed co-localization of C3d with IgG and IgM within vessels (Figure 3C), consistent with a role of IgM and IgG in complement activation. This vascular phase was followed by a second phase where C3d appeared in the parenchyma, initially perivascular at 2 hours, and more widespread by 4.5 hours post-stroke (Figure 3A). This was accompanied by a corresponding increase in parenchymal IgG and IgM signals. Notably, the initial parenchymal signals at 2h post stroke were concentrated around C3d-positive vessels (Figure 3A; higher magnification in Figure 3D), suggesting that complement activation within the vasculature leads to blood-brain barrier disruption and vessel leakage. Significantly, the escalation in C3d was at a time point after glucose levels had normalized (Figure 1B), implying that correcting glycemia alone cannot stop the cascade once it has begun.

These results indicate that hyperglycemic stress results in extremely rapid complement activation in the ischemic brain vasculature, starting within minutes of stroke onset. This is subsequently followed by severe BBB breakdown and parenchymal complement activation.

### Reperfusion Accelerates Vascular Complement Activation and Immune Protein Leakage After Hyperglycemic Stroke

The transition from vascular-confined complement activation to parenchymal propagation occurred after reperfusion (Figure 3), suggesting that reperfusion may accelerate this progression. To test this, we compared hyperglycemic stroke mice subjected to transient MCAO (with reperfusion) versus permanent MCAO (without reperfusion). As expected, at 4.5 hours post-stroke, mice that underwent reperfusion exhibited marked accumulation of IgG, IgM, and the complement activation product C3d throughout the ischemic parenchyma (Figure 4 A). In marked contrast, mice without reperfusion exhibited only minimal immune signals, largely confined to the vascular compartment. Quantification confirmed reperfusion significantly elevated levels of all three markers versus the group without reperfusion (Figure 4B). These data emphasize that reperfusion significantly amplifies vascular complement activation and BBB breakdown in hyperglycemic stroke, accelerating complement activation in the brain parenchyma.

**Figure 4.**
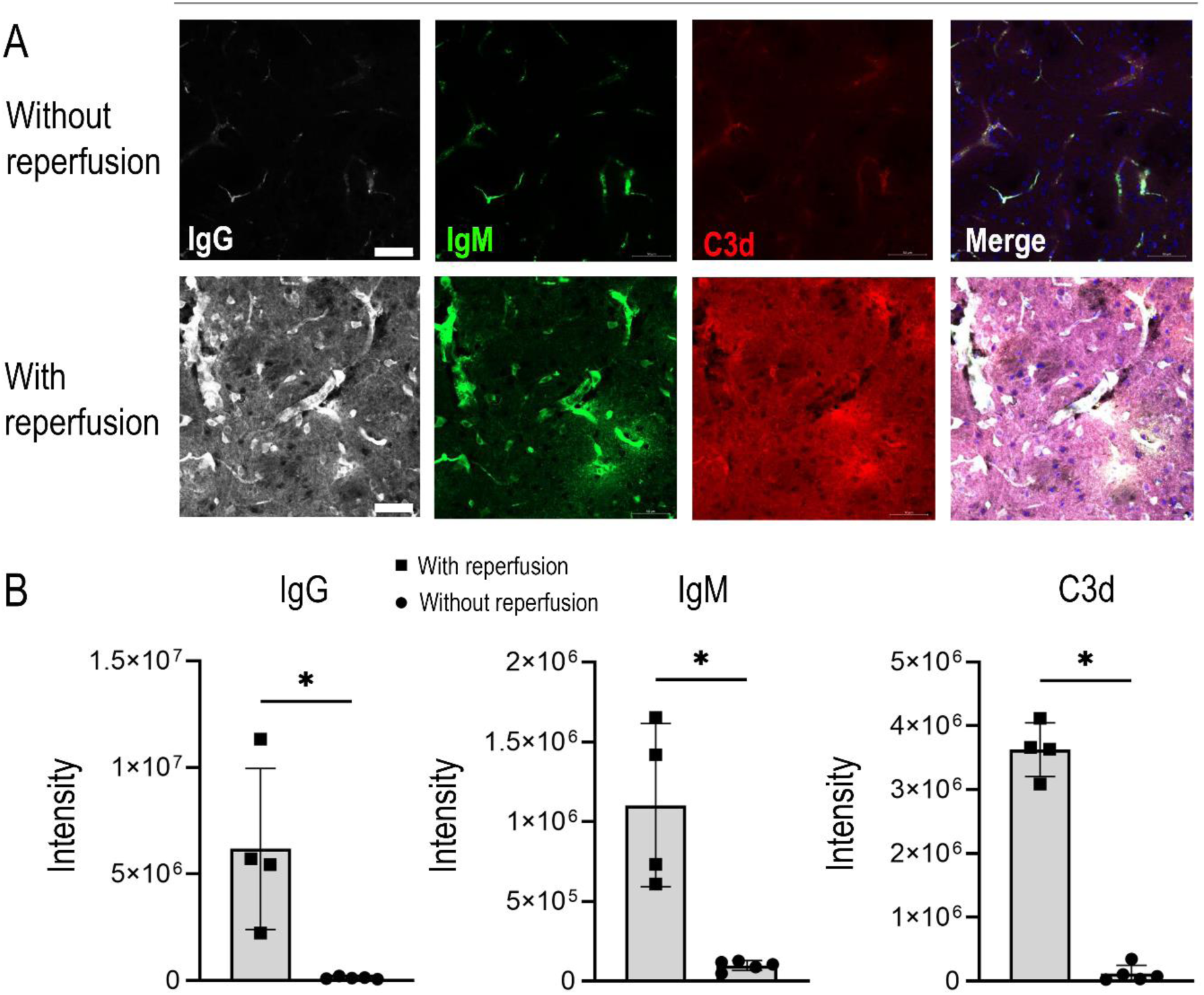
Reperfusion amplifies complement activation and BBB leakage in hyperglycemic stroke. **A-B**, Representative immunofluorescence images and quantification of IgM, IgG, and C3d at 4.5 hours post-stroke in hyperglycemic mice, with and without reperfusion. Scale bars: 50 µm. Data are presented as mean ± SD; *p < 0.05, unpaired t-test or Mann-Whitney test.

### Complement C3 Drives Vascular Immune Injury in Hyperglycemic Stroke

To determine whether complement C3 mediates the vascular disruption and poor outcomes observed in hyperglycemic stroke, we evaluated its role using both genetic and pharmacological approaches (Figure 5A, H). C3 knockout (C3 KO) mice subjected to hyperglycemic stroke showed markedly reduced staining of C3d, IgM, and IgG in the ischemic hemisphere at 24 hours post-stroke compared to wild-type (WT) controls (Figure 5B–C). Furthermore, C3 KO mice exhibited reduced neurological deficits at 24 hours (Figure 5D) and significantly better survival rates by day 5 compared to WT controls (Figure 5E). Open field behavioral tests showed that C3 KO mice exhibited improved motor and exploratory activity, evidenced by increased travel distance and center crossings at post-stroke day 5 compared to WT mice (Figure 5F-G). Notably, cresyl violet staining showed no difference in infarct size between WT and C3 KO mice at 24h post-stroke (n=6; Figure S4A–B), suggesting that the observed reductions in leakage and improved neurological function were not due to smaller infarcts.

**Figure 5.**
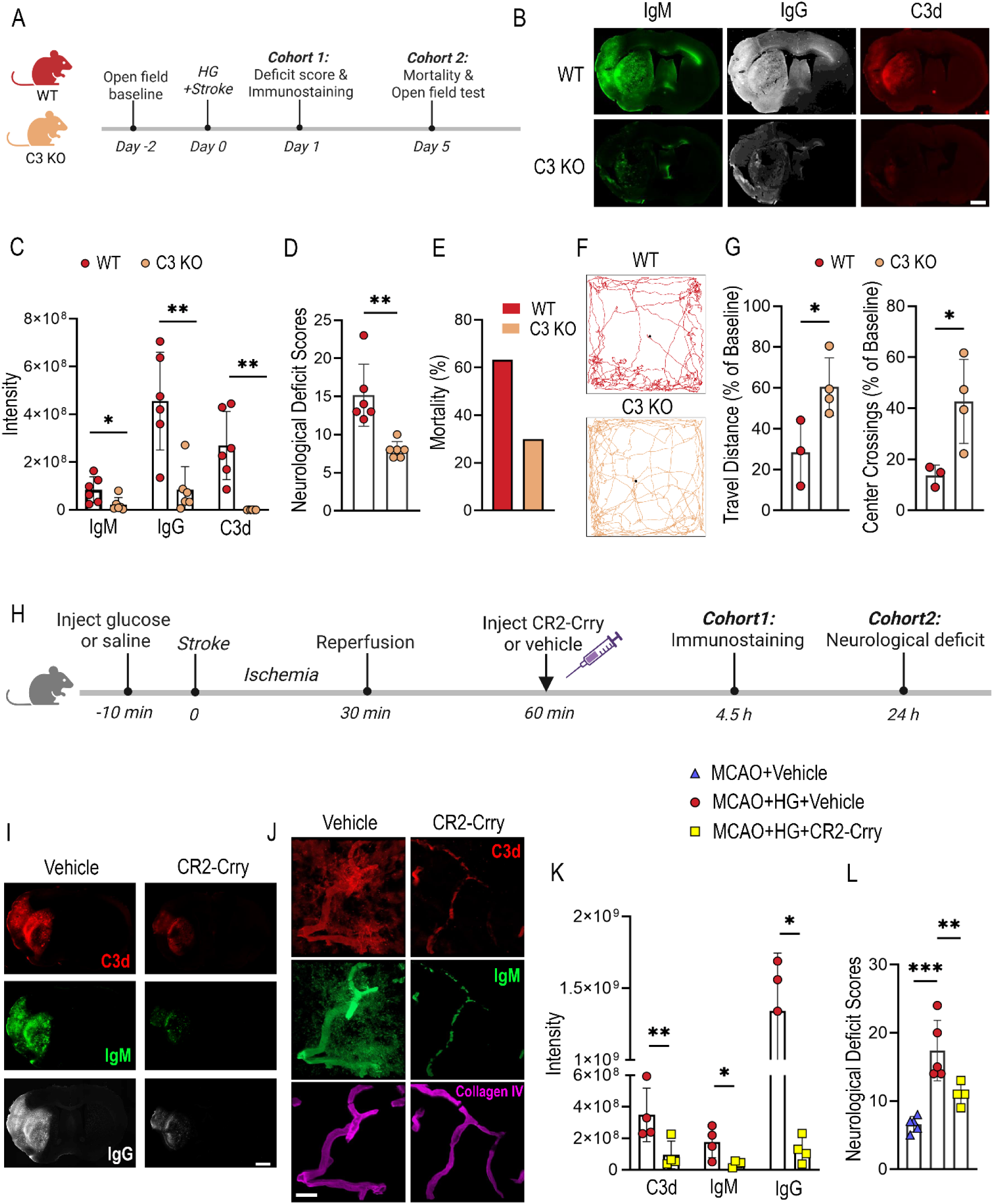
Genetic deletion or pharmacologic inhibition of C3 reduces BBB injury and improves stroke outcomes in hyperglycemic mice. **A**, Schematic timeline for two cohorts: Cohort 1 for survival and open field analysis; Cohort 2 for neurological deficit and mechanistic analysis. **B-C,** Representative whole-brain immunostaining (**B**) and quantification (**C**) of IgM, IgG, and C3d deposition in wild-type (WT) and C3 knockout (C3 KO) mice at 24 hours post-stroke. n=6/group. unpaired t-test, *p<0.05, **p<0.01. Scale bar:1000μm. **D**, Neurological deficit scores at 24 hours post-stroke. n=6/group, Unpaired t-test, **p<0.01. **E**, C3 knockout (KO) mice exhibited significantly improved survival by day 5 after hyperglycemic stroke compared to wild-type (WT) mice. Fisher-exact test, p<0.05. n=19 for WT group, n=11 for C3 KO group. **F-G**, Open field test results on day 5. **F**, Representative movement tracing map; **G**, Total travel distance and number of center crossings. n=3-4/group; unpaired t-test, *p<0.05. **H**, Schematic showing timing of CR2-Crry administration (30 minutes post-reperfusion, equal to 60 minutes post-stroke), mimicking post-thrombectomy treatment. **I and K**, Representative immunostaining (**I**) and quantification (**K**) of C3d, IgM, and IgG deposition at 4.5 hours post-stroke in MCAO + HG mice treated with CR2-Crry or vehicle. n = 4/group; unpaired t-test. *p<0.05, **p<0.01. Scale bar:1000μm. **J**, Representative immunostaining showing that CR2-Crry treatment prevented the propagation of C3 activation (C3d) and IgM deposition into the brain parenchyma. Collagen IV: vessel marker. Scale bar: 20μm. **L**, Neurological deficits at 24 hours. n=5/group. Unpaired t-test, **p<0.01; ***p<0.001. All data are represented as mean ± SD.

To explore translational potential, we tested pharmacological inhibition of C3 using CR2- Crry, a site-targeted complement inhibitor that binds activated C3 fragments^28^. CR2-Crry was administered systemically (i.p.) 30 minutes after reperfusion (Figure 5H), simulating a post-thrombectomy treatment window, during a critical period characterized by moderate vascular C3d deposition but preceding complement activation in the brain (Figure 3A). CR2-Crry-treated mice, compared to the vehicle-treated group, exhibited significantly greater IB4⁺ microvascular staining (Figure S4C-D) and reduced C3d, IgM, and IgG staining in the ischemic hemisphere at 4.5 hours post-stroke (Figure 5I–K). CR2-Crry effectively blocked the widespread complement activation and IgM/IgG deposition across brain regions (Figure S4E). Critically, targeting vascular complement at reperfusion blocked its propagation into the brain parenchyma (Figure 5J). CR2-Crry treatment also significantly improved neurological outcomes at 24 hours (Figure 5L).

Collectively, these findings highlight vascular C3 activation as a key early driver of BBB injury and exacerbated outcomes in hyperglycemic stroke, and identify a clinically actionable window following reperfusion, for targeted C3 inhibition.

### Human Stroke Brains Reveal Endothelial Glycocalyx Loss and Vascular Complement Activation

To determine whether our findings from the rodent model have translational relevance, we analyzed post-mortem brain tissues from human stroke patients and controls. Demographic and clinical characteristics of the human stroke and control samples are summarized in Table S1. We performed immunostaining for C3d, UEA I (Ulex europaeus agglutinin I, a lectin that can bind to the endothelial luminal glycocalyx), and IgG. Our confocal imaging of collagen IV and UEA I confirms the luminal localization of UEA I in human brain vessels, supporting its use as a marker of the endothelial luminal surface (Figure S5A). Representative images (at low and high magnification) revealed similar findings to our rodent model with increased vascular C3d deposition, decreased UEA I coverage, and elevated vascular and parenchymal IgG in stroke samples compared to controls (Figures 6A–C). Quantification confirmed a significant increase in IgG intensity (p < 0.05), while C3d intensity and UEA I coverage showed non-significant trends (data not shown), likely due to limited sample size.

**Figure 6.**
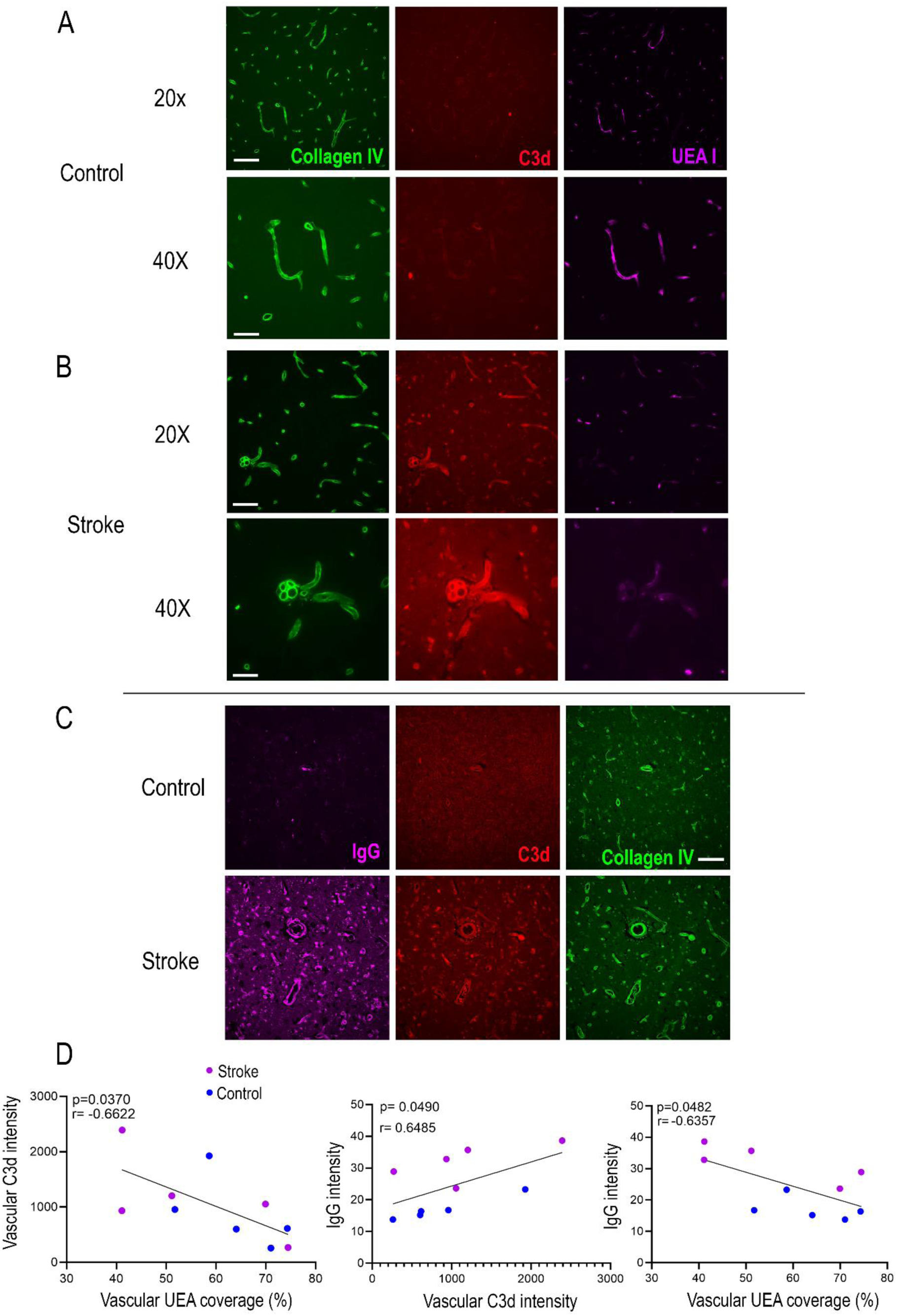
Human stroke brains exhibit endothelial glycocalyx loss and vascular complement activation associated with blood-brain barrier disruption. **A–B**, Representative confocal images of post-mortem brain tissue from control and stroke patients, stained for C3d, UEA I, and collagen IV. Scale bars: 100 μm (20 x images) and 50 μm (40 x images). n=5/group. **C**, Representative confocal images of brain tissue from control and stroke patients, stained for C3d, IgG, and collagen IV. Scale bars: 100 μm. n=5/group. **D**, Correlation analyses across stroke and control samples (n = 5/group). Left: Vascular C3d intensity negatively correlated with UEA I coverage. Middle: IgG intensity positively correlated with vascular C3d intensity. Right: IgG intensity negatively correlated with UEA I coverage. Each dot represents an individual brain; Data were analyzed using Pearson correlation.

Correlation analysis revealed that vascular C3d intensity negatively correlated with UEA coverage, and positively with IgG intensity. In parallel, loss of UEA I also correlated with increased IgG intensity (Figure 6D), suggesting that vascular complement activation and endothelial glycocalyx damage are associated—in the human stroke brain. To validate this relationship at higher resolution, we performed per-vessel correlation analysis in three independent hyperglycemic stroke brains. In all cases, we observed consistent and statistically significant negative correlations between vascular C3d intensity and UEA I coverage (Figure S5B).

Together, these findings provide the first evidence, to our knowledge, that the human stroke brain exhibits endothelial luminal surface injury and vascular complement activation, and that these vascular changes are closely interconnected, offering key mechanistic insights. Notably, all stroke samples analyzed in this study were from hyperglycemic patients, linking our observations to the hyperglycemia-associated vascular injury observed in the rodent model. These results suggest the translational relevance of our preclinical findings and highlight glycocalyx disruption and vascular complement activation as potential therapeutic targets for preserving BBB integrity after stroke.

### Pre-Thrombectomy Circulating Complement Activation Predicts Clinical Outcomes

Our identification of luminal complement activation is uniquely suited for biomarker-guided therapy as its activation yields both surface-bound (e.g. C3d) and soluble fragments (e.g., C3a, C4a, Ba) that enter the circulation (Figure S6), enabling non-invasive tracking of complement-mediated injury. We strategically leveraged this important biological feature to test whether these soluble complement activation products could serve as prognostic biomarkers in stroke patients undergoing mechanical thrombectomy. This patient population was selected based on our preclinical findings that reperfusion amplifies complement-mediated damage, and because no prior studies have evaluated the predictive value of complement in this specific clinical context.

Arterial cerebral blood was taken locally during thrombectomy, from a site proximal to the clot^29^. This sampling approach uniquely captures complement activity within the ischemic vascular microenvironment immediately prior to reperfusion, potentially increasing the sensitivity and clinical relevance of biomarker detection. The complement protein signature of the blood samples was determined using the MicroVue™ Complement Multiplex assay. A total of 66 acute ischemic stroke patients were included in the study, with key demographics summarized in Figure 7B, and additional demographic data provided in Figure S7A. Only patients achieving successful recanalization (TICI 2b–3) were included to mitigate confounds related to reperfusion variability. The range of plasma complement proteins detected in the cohort is illustrated in Figure S7B.

**Figure 7.**
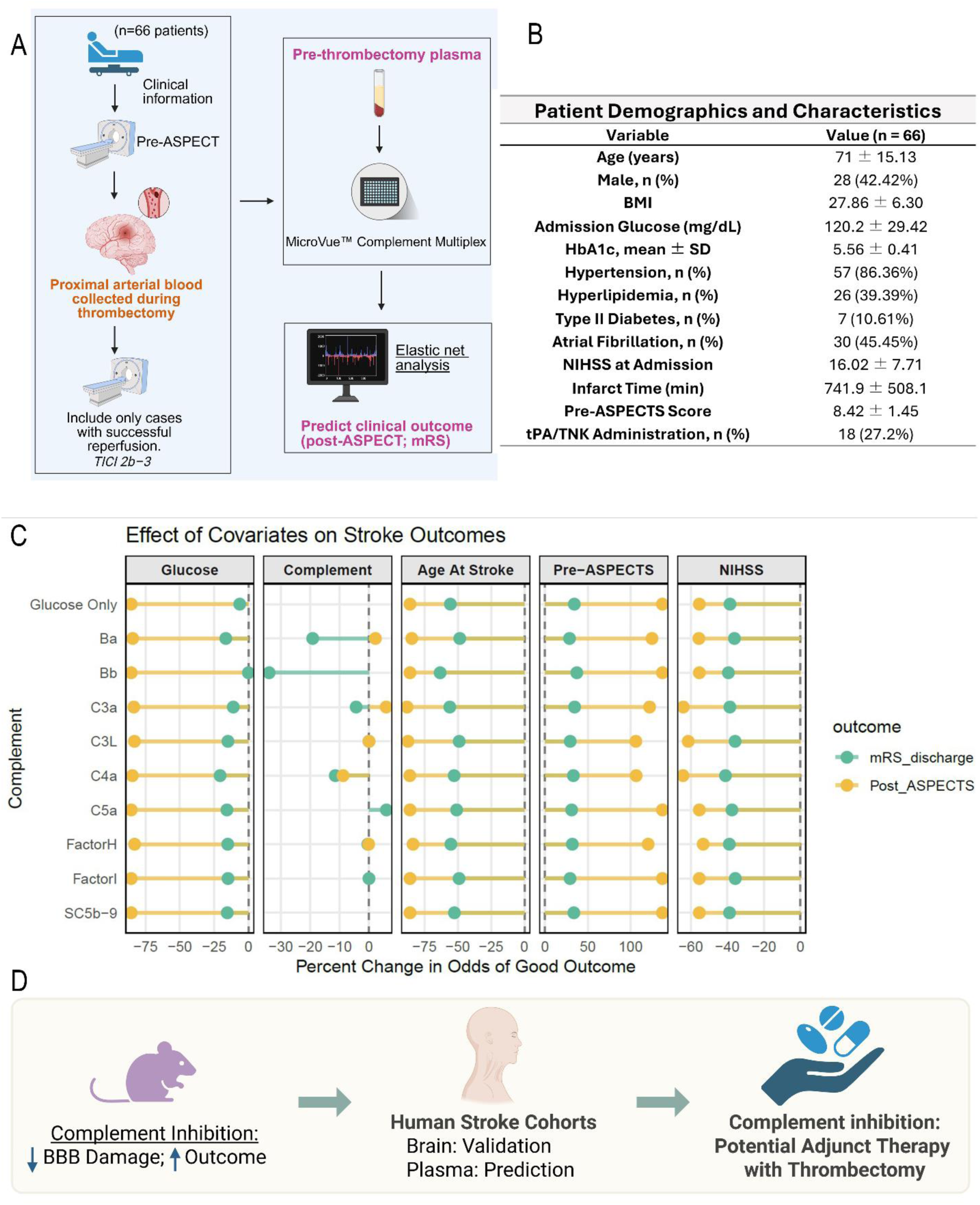
Pre-thrombectomy circulating complement activation products and blood glucose levels predict post-thrombectomy outcome. **A,** Workflow for pre-thrombectomy complement assay and outcome prediction in stroke patients. Clinical data and pre-thrombectomy blood samples were collected from ischemic stroke patients (n=66) undergoing mechanical thrombectomy. All included patients achieved good reperfusion (TICI 2b–3). EDTA plasma was processed and analyzed at National Jewish Health using the MicroVue™ Complement Multiplex assay. Complement levels and clinical information were included in unsupervised elastic net regression to predict post- ASPECT score and mRS at discharge. **B**, Patient demographics and clinical characteristics of the thrombectomy cohort (n = 66). Values are presented as mean ± SD or n (%). See Figure S7A for more information. **C**, Percent change in odds of good outcome for key predictors across stroke outcomes. Y-axis shows different complement models (with “Glucose Only” as baseline model, which include clinical parameters). X- axis represents the percent change in odds per one-unit increase in each predictor. Colors indicate outcome measures: mRS_discharge (green) and Post-ASPECTS (yellow). Panels display effects of Glucose, specific Complement factors, Age, pre- ASPECTS score, and NIHSS. Negative values indicate decreased odds, positive values indicate increased odds of good outcomes. Dashed vertical lines represent no effect. See Table S2 for full covariate results. **D**, Schematic summary of the translational relevance of this study, illustrating mouse-to-human insights.

To identify independent predictors of outcome we performed elastic net regression with 5-fold cross-validation—a robust regularization approach that enhances generalizability. Clinical data—including NIHSS scores, pre-ASPECT imaging, admission glucose levels, and thrombolytic use (Alteplase [tPA] or Tenecteplase [TNK])—were included in the models. This approach enables data-driven selection of the most predictive features while minimizing the risk of overfitting. To determine which complement components independently predict stroke outcomes while minimizing multicollinearity, each complement protein was added individually to a base clinical model containing all recorded demographic and clinical variables (Figure 7B and Figure S7A). This allowed for the assessment of each complement protein’s independent predictive value for post- ASPECT and modified Rankin Scale (mRS) outcomes. As expected, conventional predictors such as pre-ASPECT score, age, and NIHSS were strongly associated with worse post-ASPECT scores and mRS functional outcomes; these associations remained stable when complement proteins were added individually (Figure 7C), validating the model framework. In addition, the elastic net models demonstrated strong predictive accuracy, area under the curve (AUC), sensitivity, and specificity with a minimum R^2^ of 0.803, 0.861, 0.69, and 0.853 for each parameter, respectively (Figure S7C).

Plasma complement components including Ba, Bb, C4a, and C3a, significantly predicted worse mRS scores at discharge, even accounting for clinical factors such as pre-ASPECT, age, NIHSS, glucose, and all other covariates included in the model (Figure 7C). Notably, complement protein Bb showed predictive strength comparable to Pre-ASPECT and NIHSS. While C4a, Ba, and C3a had smaller effect sizes, they exhibited a broader dynamic range across patients (Figure S7B), which could still translate into substantial differences in clinical outcomes. For example, C4a has a range of 2420 units; given an odds ratio of -11.47 per unit, this could represent a major shift in risk across patients with low versus high levels. These findings suggest that soluble complement activation products could serve as independent prognostic biomarkers of functional outcomes in thrombectomy-treated stroke patients.

Admission glucose levels (Figure 7C), which ranged from 57 to 225 mg/dL, emerged as a strong predictor of worse post-ASPECT score, with an effect size comparable to age and NIHSS admission. Notably, this suggests that a one-unit increase in glucose has a similar impact on outcome as a one-unit increase in age or NIHSS score, remarkable given that glucose values span a much broader range. This wide dynamic range implies that glucose could exert a substantial influence on acute stroke brain injury. Consistent with this, 6 of the 7 patients who died in the hospital were hyperglycemic, highlighting the acute detrimental effect of hyperglycemia on mortality after stroke.

For the functional outcome measured by mRS, the base model showed that higher glucose was associated with worse outcomes. This association persisted across all models that included complement proteins and was accompanied by an increased effect size (except in the model including Bb), suggesting that elevated glucose contributes to poor functional outcomes and that there may be an interaction between glucose and complement activation in influencing stroke outcome.

To our knowledge, this is the first study to demonstrate that pre-thrombectomy complement activation independently predicts post-thrombectomy functional recovery. This has important clinical implications, as it could help identify patients who may benefit from complement-targeted adjunct therapies (Figure 7D).

## Discussion

This study reframes hyperglycemic stroke as an acute vascular luminal problem and provides the first evidence that vascular complement activation is a key driver and therapeutic target linking acute hyperglycemia to cerebrovascular injury. Although hyperglycemia is a common metabolic stressor affecting ∼40% of stroke patients and strongly associated with poor outcomes, effective targeted interventions are lacking. In this study, we define a Metabolic–Complement–Vascular (MCV) axis, in mouse and human stroke brains, in which hyperglycemia rapidly disrupts the endothelial glycocalyx, initiating luminal immunoglobulin deposition, complement activation, and blood–brain barrier (BBB) disruption within hours of stroke onset (Figure 8). Remarkably, these vascular changes began very rapidly, within 15 minutes of stroke onset, and were intensified by reperfusion, persisting despite glucose normalization. Using both genetic and pharmacologic approaches, we demonstrate a central role for complement C3 in mediating hyperglycemia-induced BBB leakage and stroke injury. Importantly, blocking vascular complement activation at 30 min post-reperfusion significantly preserved BBB integrity and improved outcomes in hyperglycemic rodent stroke model, highlighting a clinically relevant therapeutic window. We also provide the first clinical evidence that pre-thrombectomy plasma levels of complement activation products predicted post-thrombectomy outcomes in stroke patients, supporting the potential for pre-intervention risk stratification. These findings establish vascular complement as both a mechanistic driver and a clinically actionable target in hyperglycemic stroke, highlighting vascular complement as a promising adjunctive target at the time of thrombectomy.

**Figure 8.**
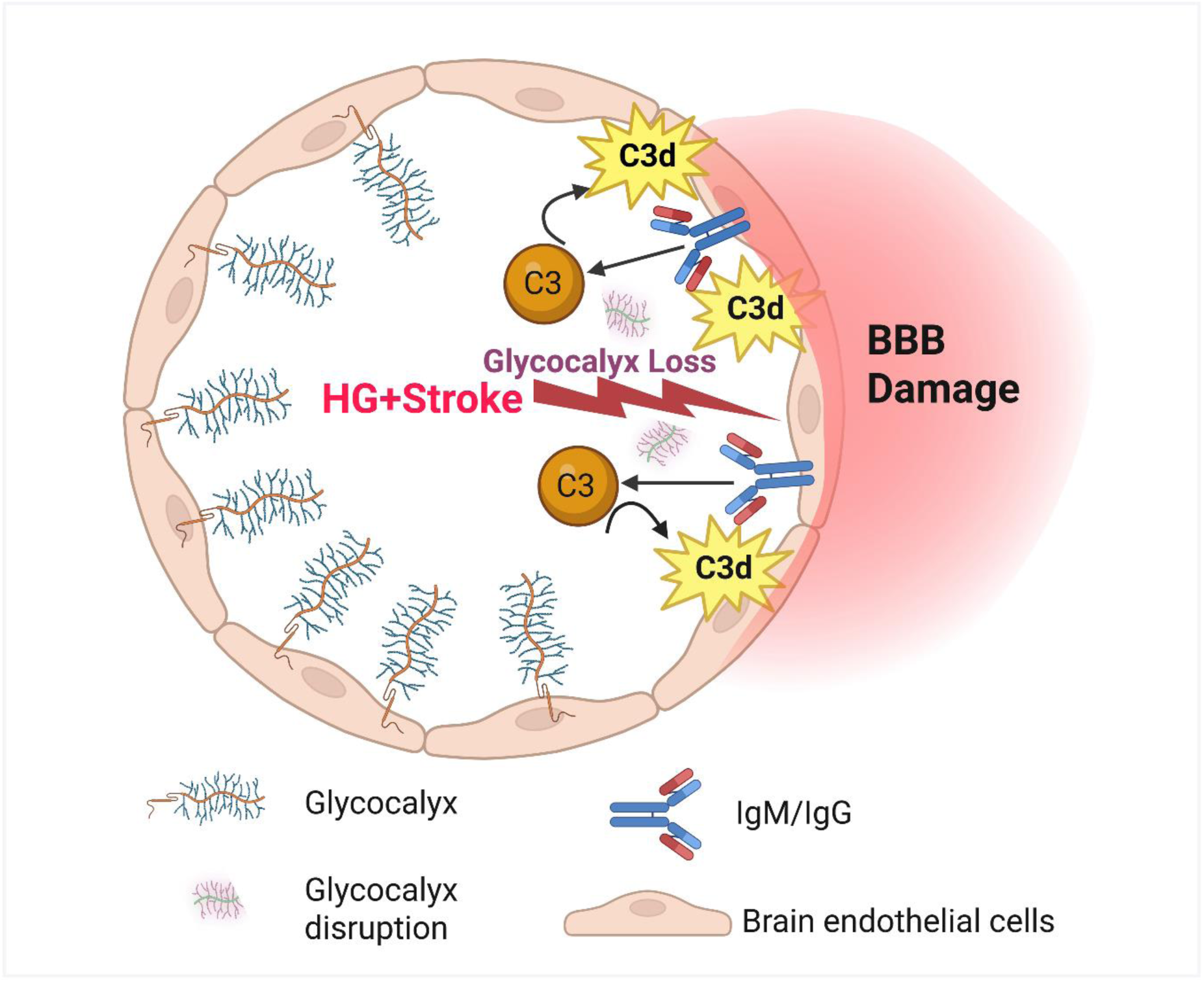
Schematic model of the Metabolic–Complement–Vascular (MCV) axis linking hyperglycemia stress to luminal damage and blood-brain barrier (BBB) breakdown. In this model, hyperglycemia (HG) with stroke induces rapid disruption of the endothelial glycocalyx, exposing the luminal interface to IgM/IgG accumulation, which can trigger vascular complement activation (C3 → C3d). This luminal-initiated cascade drives blood–brain barrier (BBB) leakage and subsequent injury. The sequence highlights a novel mechanism through which vascular complement becomes an actionable target for early intervention in hyperglycemic stroke. Reperfusion significantly exacerbates this response and sets the vascular clock for timely therapeutic targeting.

The luminal glycocalyx, rich in glycoproteins and proteoglycans, plays a crucial role in maintaining vascular integrity and endothelial function^15^. Our discovery of rapid glycocalyx degradation in mice stroke brains following acute hyperglycemia offers a novel insight into the timing and pathophysiology of hyperglycemic stroke. Significantly, we present the first evidence in human stroke brain tissue of endothelial luminal glycocalyx loss, suggesting the translational relevance of our findings. Previous hyperglycemic stroke studies focused on oxidative stress and leukocyte-endothelial adhesion in the context of BBB disruption. These phenomena are potentially linked with glycocalyx breakdown as oxygen radicals can degrade components of the glycocalyx^30^, and leukocyte adhesion may be more likely in the absence of the protective luminal glycocalyx barrier. While two recent studies reported glycocalyx loss in stroke^31,32^ and its association with BBB compromise, our data highlight hyperglycemia as a potent accelerator of glycocalyx degradation. In our model, just a brief period of hyperglycemia (less than 2 hours) resulted in almost complete loss of glycocalyx in the ischemic territory within 4.5 hours of stroke, contrasting with the mostly intact glycocalyx in normoglycemic stroke. This underscores the metabolic vulnerability of the brain endothelial glycocalyx to high glucose levels, which is further supported by studies in the periphery demonstrating the impact of hyperglycemia on glycocalyx shedding^17^. However, unlike peripheral vessels where hyperglycemia alone damages the glycocalyx, cerebral vessels did not show glycocalyx loss beyond the ischemic region, emphasizing that cerebral vascular injury required the combined stress of hyperglycemia plus ischemia. This is consistent with other studies reporting that the glycocalyx in cerebral vessels is much more robust and resistant to injury than its peripheral counterpart^18,19^. This double-hit model characterizes hyperglycemic stroke as an aggressive form of vascular injury, distinct from the milder BBB disruption seen in normoglycemic stroke. These findings establish glycocalyx as a specific site of vascular vulnerability and a structural link connecting systemic metabolic disturbances to cerebral neurovascular injury. The results offer the potential for developing new strategies to monitor and protect endothelial luminal health in stroke and other brain diseases driven by metabolic stressors.

Our histological data from mouse and human brain tissue indicate an association between glycocalyx damage and vascular complement activation, suggestive of a model in which glycocalyx breakdown facilitates luminal binding of the complement activators IgG and IgM, possibly through exposure of neoepitopes on endothelial cells^33^, leading to luminal complement C3 activation. Our model reframes early stroke injury from being solely a brain-centric process to one that includes a metabolic–complement axis that initiates early and critical pathological events at the luminal surface of the cerebrovascular interface involving early glycocalyx disruption. Previous studies of complement in stroke have focused primarily on parenchymal complement activation leading to neuronal injury^34^, with only sporadic reports of vascular involvement^33,35,36^, which focused more on the concept that activation of complement in the vasculature impairs the recovery of blood flow^35^.

Building upon this, we propose a mechanistic framework—the Metabolic–Complement– Vascular (MCV) axis—in which systemic hyperglycemic stress, in the context of ischemic stroke, triggers glycocalyx loss and complement activation within the disrupted vessel lumen. This complement activation then propagates from the lumen to perivascular spaces and into the brain parenchyma, where systemic C3 influx and local production of C3 by brain-resident or infiltrating immune cells can contribute to parenchymal complement activation^37,38^. The MCV axis thus connects metabolic stress to subsequent brain injury via complement-mediated damage at the vascular interface, integrating metabolic, vascular, and immune aspects in stroke pathophysiology. Critically, reperfusion accelerates this axis: restoring blood flow amplifies complement activation and immune proteins propagation into the brain. This is consistent with prior studies where systemic complement depletion conferred benefit only in transient (reperfusion) stroke models^39,40^. These findings expose a fleeting, but actionable therapeutic window immediately following reperfusion.

Our study reveals translational opportunities. Despite advances in reperfusion therapies, hyperglycemia remains a major determinant of poor outcomes ^25^, a fact further supported by our unbiased elastic net regression models. This is significant as nearly 40% of stroke patients present with acute hyperglycemia, whether this be from stress hyperglycemia^25^ or diabetes, and no targeted therapies exist. Insulin therapy alone provided limited benefit to patients^8^ and our data suggest this is because hyperglycemia acts as a rapid trigger initiating a cascade of harmful events—the MCV axis—in the ischemic vasculature within minutes of stroke onset. Critically, we showed that glucose normalization fails to halt vascular complement activation and parenchymal progression once the MCV axis is triggered. In essence, by the time the patient gets to the hospital it is too late to target glucose. However, we demonstrate that early complement inhibition post-reperfusion shows promise in maintaining blood-brain barrier integrity, restricting complement spread into the brain, and enhancing neurological function. This offers a systemically deliverable, luminal-targeted, and time-defined therapeutic strategy for hyperglycemic stroke. The translational potential is strengthened by our demonstration of vascular complement activation in stroke patients. Importantly, our findings redefine complement-targeting strategies in the CNS: rather than overcoming the blood–brain barrier to reach brain parenchyma—a major obstacle^41^—we show that pathogenic complement activation occurs at the vascular luminal surface in hyperglycemic stroke, exposing an accessible target for systemic intervention.

Intervention timing is crucial: vessel reopening accelerates vascular complement overactivation and its propagation into the brain, exposing a brief but critical post-reperfusion window where vascular immune injury can be intercepted, particularly in metabolically vulnerable patients. This adds to the traditional concept in stroke management of “time is brain,” emphasizing that reperfusion also sets a vascular clock. Importantly, the luminal complement perspective offers a distinct biological advantage: its activation generates both surface-bound effectors and soluble fragments that release into circulation, making otherwise invisible luminal vascular injury trackable in plasma.

This supports the potential of biomarker-guided, tailored adjunct therapy—a feature not typically feasible with brain-targeted interventions. Our clinical plasma data reinforces this approach, showing that pre-thrombectomy C3-activation markers predict post-thrombectomy outcomes independently, demonstrating the potential of biomarker-guided complement inhibition as a viable adjunct strategy during reperfusion.

### Limitations and Future Studies

While our study provides strong mechanism and translational evidence, the clinical cohort is modest in size and drawn from a single high-risk geographic region. Prospective, larger-scale studies are needed to validate C3 activation as a predictive biomarker and to assess the efficacy of complement-targeted therapies in stroke. A larger cohort will also enable comparison of diabetic and non-diabetic patients to determine if the two groups have a similar response to hyperglycemic stroke; our current cohort predominantly consists of non-diabetic patients aligning with our preclinical model. While preclinical experiments used males to minimize variability, our human samples incorporated both sexes of varying ages, supporting broader relevance. Although we did not delineate the precise pathways driving vascular complement activation, the association of Bb, Ba (alternative pathway) and C4a (classical/lectin pathway) with clinical outcomes suggests involvement of multiple arms of the cascade. Furthermore, whether dual targeting of glycocalyx preservation and complement inhibition yields superior outcomes remains an important avenue for future research.

In conclusion, our study highlights the critical role of the vascular luminal glycocalyx and vasculature complement activation in hyperglycemic stroke and identifies a Metabolic–Complement–Vascular (MCV) axis that links systemic metabolic stress to glycocalyx disruption, vascular complement activation, and progressive neurovascular injury and BBB breakdown. Our data provides a new understanding of hyperglycemic stroke pathophysiology and also proposes concrete clinical applications, such as complement inhibition as an adjunct therapy during reperfusion, and the use of plasma complement activation products for potential patient stratification. These insights could refocus therapeutic efforts to protect the cerebrovascular interface, particularly from the luminal side, including preserving glycocalyx integrity and preventing complement activation. Beyond stroke, the MCV axis may represent a broader paradigm and therapeutic target across metabolically driven cerebrovascular disorders.

## Methods

### Animal Experiments

All animal procedures were approved by the Stanford University Institutional Animal Care and Use Committee and performed in accordance with the National Institutes of Health (NIH) Guidelines for the Care and Use of Laboratory Animals. To avoid hormonal variability in infarct size associated with the estrous cycle, we used male C57BL/6 mice (10–11 weeks old). Mice were obtained from The Jackson Laboratory and housed in a controlled environment with a 12-hour light–dark cycle, constant temperature and humidity, and ad libitum access to standard chow and water. All animals underwent a one-week acclimatization period prior to experimental procedures. Animals were randomly assigned to experimental groups. Sample sizes were based on prior studies and established practices in the field. All efforts were made to minimize animal suffering and reduce the number of animals used. Randomization and blinding were applied throughout the study, including during surgery, treatment allocation, outcome assessment, and data analysis.

### MCAO Stroke Model and Induction of Acute Hyperglycemia

For MCAO, mice were anesthetized with 5% isoflurane for induction (5 minutes), followed by maintenance with 2% isoflurane using a vaporizer (Vetequip). Oxygen was delivered as a mixture of air and oxygen at a 4:1 ratio. Body temperature was maintained at 36.5–37°C throughout the procedure using a feedback-regulated DC temperature controller (FHC, 40-90-8D). Heart rate and respiration were assessed at 15-minute intervals to ensure physiological stability. To induce acute hyperglycemia, glucose was injected (2.2 g/kg, i.p.) at 10 min before stroke ^24^. The same volume of saline injection was used as a control. A midline neck incision was made to expose the common carotid artery (CCA), external carotid artery (ECA), and internal carotid artery (ICA). A silicone-coated monofilament (Doccol, #702045PK5Re, 0.20 ± 0.01 mm coating diameter) was inserted via the ICA and advanced to occlude the origin of the MCA. After 30 minutes of occlusion, the filament was gently withdrawn to initiate reperfusion, mimicking the clinical thrombectomy scenario. The incision was then closed using sterile sutures. Postoperatively, mice received subcutaneous buprenorphine (0.1 mg/kg) for analgesia and 0.5 ml sterile saline (Hospira, NDC 0409-4888-10, i.p.) to prevent dehydration. Mice were allowed to recover in a warm recovery cage before being returned to their home cages. For subacute survival studies, mice received 1 ml of sterile saline (subcutaneous) twice daily for 7 days and were provided with a nutrient-enriched diet gel to support recovery. Body weight, general status, and survival were monitored daily.

### Transgenic Mice

To investigate the role of complement C3 in hyperglycemia-exacerbated stroke pathophysiology, we utilized C3-deficient (C3⁻/⁻) mice on a C57BL/6J background (B6;129S4-C3tm1Crr/J, Stock No. 003641, The Jackson Laboratory). Age-matched male wild-type (WT) C57BL/6J mice (10-11 weeks old) were used as controls. Both C3⁻/⁻ (C3 KO) and WT mice were subjected to identical MCAO and hyperglycemia protocols as described above. All surgical, post-operative, and outcome assessments were performed in a blinded manner.

### CR2-Crry Administration

To evaluate the therapeutic potential of targeted complement inhibition, we utilized CR2- Crry, a site-specific complement inhibitor kindly provided by Dr. Stephen Tomlinson (Medical University of South Carolina). CR2-Crry is a recombinant fusion protein comprising the four N-terminal short consensus repeat (SCR) domains of complement receptor 2 (CR2), which mediate binding to sites of complement activation, linked to the inhibitory domain of murine Crry, a potent regulator of C3 activation^21^. CR2-Crry was administered intraperitoneally at a dose of 0.5 mg per mouse^21^ (in 100 µL chilled calcium- and magnesium-free PBS), 30 minutes after reperfusion. Control mice received an equivalent volume of vehicle (chilled PBS without calcium or magnesium). All injections were performed using insulin syringes under sterile conditions. Investigators were blinded to treatment groups during administration and subsequent analysis.

### Evans Blue Assay

To assess blood–brain barrier (BBB) permeability, Evans blue (EB; Sigma-Aldrich) dye—an albumin-binding tracer that normally does not cross the intact BBB—was used. Two hours after MCAO induction, mice (including both stroke and sham-operated groups) received an intravenous injection of 2% EB solution in saline (4 mL/kg) via the femoral vein. At 2.5 hours post-injection, mice were transcardially perfused with 35 mL phosphate-buffered saline (PBS) to remove intravascular EB. Following perfusion, brain hemispheres and liver tissues were dissected, weighed, and homogenized in N,N-dimethylformamide (Sigma-Aldrich). Samples were incubated overnight at 55 °C and then centrifuged at high speed. When needed, supernatants were diluted to ensure measurements fell within the linear range of the standard curve. Optical density (OD) of the supernatants was measured at 620 nm using a microplate reader (Tecan Infinite M200 Pro). EB concentrations were calculated based on a standard curve generated using serial dilutions of known EB concentrations. Liver Evans blue levels were quantified in parallel to confirm consistent systemic delivery across experimental groups. No significant differences in liver EB content were observed between groups (Figure S2B), supporting the validity of brain EB measurements.

### Hemorrhagic Transformation and Brain Swelling

Hemorrhagic transformation (HT) was evaluated using both microscopic and macroscopic approaches. For microscopic assessment, brain sections were stained for hemoglobin, and the number of parenchymal hemoglobin-positive clusters was manually counted across 10 coronal slices per brain. The cumulative number of clusters was recorded as the microscopic HT burden. For macroscopic assessment, post- mortem brain slices were visually inspected for the presence of hemorrhage. HT was defined as clearly visible intracerebral blood accumulation within the infarcted tissue. Each brain was sectioned into 7 coronal slices, and the number of slices exhibiting visible hemorrhage was quantified per animal to determine the extent of macroscopic HT.

Brain swelling was assessed by measuring hemispheric enlargement using ImageJ software (NIH). The swelling ratio was calculated as the area of the ipsilateral (ischemic) hemisphere divided by the area of the contralateral (non-ischemic) hemisphere on matched coronal sections^42^.

### Post-mortem Verification of Brain Infarction by TTC Staining

For mice that died during the 14-day post-stroke observation period (Fig 1), brain infarction was verified post-mortem using 2,3,5-triphenyltetrazolium chloride (TTC) staining. Brains were harvested, sectioned into 1-mm coronal slices using a brain matrix, and incubated in 2% TTC (Sigma-Aldrich) prepared in phosphate-buffered saline for 30 minutes in the dark. Infarcted tissue remained pale, whereas viable tissue stained red. The presence of infarction was required for inclusion in outcome analysis. Infarction was confirmed in all mice that died during this period.

### Cresyl Violet Staining and Infarct Volume Analysis

To assess infarct size at the acute phase after stroke, mice were sacrificed at 24 hours post-reperfusion. Brains were harvested, fixed in 4% paraformaldehyde overnight at 4°C, and then cryoprotected in 30% sucrose before sectioning. Coronal brain sections (30– 40 μm thick) were collected and mounted onto Superfrost Plus glass slides (Thermo Fisher Scientific), then air-dried overnight at 4°C. Slides were stained with 0.1% cresyl violet (Nissl stain) in distilled water for 10–15 minutes, followed by sequential dehydration through graded ethanol (70%, 95%, 100%), clearing in xylene, and cover- slipping with mounting medium. Infarct areas were visualized under a light microscope and quantified using ImageJ software (NIH).

Images of stained sections were scanned, and infarct analysis was performed using ImageJ (NIH). To correct for potential brain edema and avoid underestimation of infarct size, infarct volume was calculated by comparing the area of intact (non-infarcted) tissue in the ipsilateral (stroke-affected) hemisphere to the total area of the contralateral hemisphere ^43,44^. For each of 10 coronal slices: Infarct area = Contralateral hemisphere area – Ipsilateral non-infarcted area. The infarct areas from all slices were summed to obtain the total infarct volume. To express infarct size as a percentage, total infarct volume was divided by the summed contralateral hemisphere volume and multiplied by 100: Percent infarct = (Infarct volume / Total contralateral volume) × 100. Mice without evidence of brain infarction were pre-specified for exclusion; however, all mice exhibited infarcts at the designated acute phase time points.

### Immunostaining

Mice were transcardially perfused with 40 mL of phosphate-buffered saline (PBS) to remove circulating blood, followed by 3% paraformaldehyde (PFA) in PBS to fix brain tissue. Brains were post-fixed overnight in 3% PFA with 15% sucrose, then snap-frozen and stored at −80 °C until sectioning. Coronal brain sections were cut at 30 μm thickness using a microtome and stored as free-floating sections in antifreeze medium (30% ethylene glycol, 30% glycerol in PBS) at −20 °C.

For immunostaining, sections were washed in PBS (3×) and incubated in pre-heated antigen retrieval buffer (0.5 M sodium citrate with 0.05% Tween-20 in deionized water) at 60 °C for 20 minutes. After cooling, sections were blocked for 1 hour at room temperature in blocking solution containing 5% donkey serum and 10% bovine serum albumin (BSA) in PBS with 0.03% Triton X-100. Primary antibodies used included: anti-IgM (Jackson ImmunoResearch, 715-545-140), anti-IgG (Jackson ImmunoResearch, 715-605-151), anti-C3d (R&D Systems, AF2655)^45^, anti-CD31 (BioLegend, 102516), anti-hemoglobin (Abcam, ab92492), anti-collagen IV, anti-GFAP (Abcam Ab4674), anti- CD68 (Ab53444), and IB4 lectin (Invitrogen, I21413). Tissues were incubated overnight at 4 °C with primary antibodies diluted in blocking buffer, then washed in PBS with 0.3% Triton (3×). Secondary antibodies (donkey anti-goat, donkey anti-rabbit, donkey anti-rat) were applied for 2 hours at room temperature, followed by nuclear staining with Hoechst 33342 (Invitrogen, H3570). Sections were then mounted on glass slides with antifade mounting medium and dried overnight prior to imaging. For mice that survived until the end of the experiment (day 14), brain injury presence was confirmed by CD68/GFAP double immunostaining to validate infarct involvement for outcome inclusion. One mouse in the WT MCAO+HG group was excluded from analysis in Fig. 1A (Cohort 1), and another in Fig. 5A (Cohort 2), due to lack of infarct evidence.

### Confocal and Widefield Imaging

Brain sections processed for immunostaining were imaged using a Zeiss LSM confocal microscope for high-resolution and 3D visualization, and a Keyence BZ-X series microscope for stitched low-magnification imaging of large brain areas. For confocal imaging, Z-stack images were acquired at 1 µm intervals to capture the full depth of the section. Three-dimensional reconstruction and visualization were performed using Zeiss ZEN software. Keyence images were acquired with consistent exposure settings and stitched automatically using built-in software to generate whole-slice overviews. Imaging acquisition parameters were kept constant across experimental groups.

### Electron Microscopy for Endothelial Glycocalyx Assessment

To examine the ultrastructural integrity of the endothelial luminal glycocalyx, transmission electron microscopy (TEM) was performed on brain tissues collected 4.5 hours after stroke. Mice were transcardially perfused with sodium cacodylate buffer, followed by a fixative solution containing 2% paraformaldehyde and 2.5% glutaraldehyde in 0.1 M sodium cacodylate buffer (pH 7.4). Samples were fixed in Karnovsky’s fixative: 2% Glutaraldehyde (EMS Cat# 16000) and 4% formaldehyde (EMS Cat# 15700) in 0.1M Sodium Cacodylate (EMS Cat# 12300) pH 7.4 for 1 hr. then placed on ice 24 hours. The fix was replaced with cold/aqueous 1% Osmium tetroxide (EMS Cat# 19100) and were then allowed to warm to Room Temperature (RT) for 2 hrs rotating in a hood, washed 3X with ultrafiltered water, then en bloc stained in 1% Uranyl Acetate at RT 2hrs while rotating. Samples were then dehydrated in a series of ethanol washes for 30 minutes each @ RT, beginning at 50%, 70% EtOH then moved to 4oC overnight. They were placed in cold 95% EtOH and allowed to warm to RT, changed to 100% 2X, then Propylene Oxide (PO) for 15 min. Samples are infiltrated with EMbed- 812 resin (EMS Cat#14120) mixed 1:2, 1:1, and 2:1 with PO for 2 hrs each with leaving samples in 2:1 resin to PO overnight rotating at RT in the hood. The samples are then placed into EMbed-812 for 2 to 4 hours then placed into molds w/labels and fresh resin, orientated and placed into 65°C oven overnight. Sections were taken around 80nm using an UC7 (Leica, Wetzlar, Germany) picked up on formvar/Carbon coated 100 mesh Cu grids, stained for 40seconds in 3.5% Uranyl Acetate in 50% Acetone followed by staining in Sato’s Lead Citrate for 2 minutes. Observed in the JEOL JEM-1400 120kV. Images were taken using a Gatan OneView 4k X 4k digital camera.

### Open Field Test

Animals were placed in a 40 x 40 cm open-field arena and video-recorded at 30 frames per second for 10 minutes at baseline (pre-stroke) and at multiple post-stroke timepoints. Animal position was extracted using binary mask thresholding of the animal body against the background, and the centroid of this mask was calculated to determine animal coordinates at each frame. Total distance traveled was computed from frame-by-frame centroid displacements.

The center zone of the arena was defined as a centrally positioned square occupying 50% of the total arena area. Time spent in the center was calculated based on whether the animal’s centroid was located within this region at each frame. A center entry was defined as any transition of the centroid from outside to inside the center zone.

### Neurological Deficit Scores

Neurological function was assessed using a composite scoring system that included both general and focal deficit components^46^. Focal deficits comprised evaluations of motor function, sensory response, and reflexes. Scoring was performed by trained investigators who were blinded to experimental group assignments.

### Source of Postmortem Human Brain Tissue

Postmortem human brain tissue was sourced from the Human Brain Biorepository of Neurological Disorders at the University of Texas Health Science Center at Houston. The study was granted an exemption by the Committee for the Protection of Human Subjects at the same institution (*HSC-MS-22-0982-UTHealth Neuropathology Core*). Written consent was obtained from the next-of-kin for all patients. Table S1 provides detailed demographics and clinical characteristics of the postmortem cases included in this study. Case selection was determined based on final neuropathology reports.

### Immunofluorescence in Postmortem Human Samples

Immunofluorescence analysis was performed on formalin-fixed, paraffin-embedded human brain sections, following established protocols ^47,48^. Two sets of slides, derived from identical cases and brain regions, were used to investigate (1) the relationship between C3d and blood-brain barrier (BBB) integrity and (2) the association between C3d and endothelial cell damage. Tissue sections were deparaffinized and subjected to heat-induced antigen retrieval using citrate buffer (pH 6.0) at 99°C for 20 minutes. Subsequently, samples were blocked for 1 hour at room temperature with a blocking buffer consisting of 5% donkey normal serum, 1% bovine serum albumin, and 0.3% Triton X-100 in phosphate-buffered saline (PBS). The sections were then incubated overnight at 4°C with primary antibodies diluted in blocking buffer: mouse monoclonal anti-Collagen IV (Abcam, ab273607; 1:100) and rabbit polyclonal anti-C3d (Abcam, ab136916; 1:100). After washing with PBS to remove unbound antibodies, slides were incubated for 2 hours at room temperature with secondary antibodies: donkey anti-mouse IgG conjugated to Alexa Fluor 488 and donkey anti-rabbit IgG conjugated to Alexa Fluor 594 (both 1:200, Jackson ImmunoResearch). Additionally, for slide set 1, donkey anti-human IgG conjugated to Alexa Fluor 647 was used to assess BBB integrity, while for slide set 2, endothelial cell marker Lectin UEA1 conjugated to DyLight 647 (Vector Labs, DL-1068-1; 1:500) was applied to assess endothelial cell damage.

### Imaging and Quantification of C3d and IgG Expression in Human Brain Samples

Imaging was performed using a Leica THUNDER Imager DMi8 with a 20x objective lens. Ten images were acquired from both the infarct and peri-infarct regions of stroke patients, and ten images were captured from the occipital cortex of neurologically normal (control) individuals. The infarct and peri-infarct regions in stroke cases, as well as the cortical regions in controls, were identified using hematoxylin and eosin staining and verified by a neuropathologist. Image analysis was conducted with ImageJ software by an investigator blinded to case characteristics.

### Human Plasma Sample Collection and Clinical Data Extraction

Human plasma samples were obtained from patients undergoing cerebrovascular procedures at Albert B. Chandler Medical Center, University of Kentucky, between August 2017 and September 2023. All procedures were conducted under the Blood and Clot Thrombectomy Registry and Collaboration (BACTRAC) protocol (ClinicalTrials.gov ID NCT03153683), as previously described^29^. The study was approved by the University of Kentucky Institutional Review Board (IRB #48831). Informed consent was obtained from the patient or their Legally Authorized Representative (LAR) as soon as feasible following collection. Eligible participants were male or female patients aged ≥18 years. Exclusion criteria included pregnancy, breastfeeding, or incarceration at the time of enrollment.

Arterial blood samples were collected locally during thrombectomy, proximal to the clot, using K₂ EDTA tubes. Samples were centrifuged to isolate plasma and stored at –80 °C until analysis. Sample handling and processing followed the BACTRAC protocol in accordance with institutional biospecimen preservation standards. Clinical data were extracted from the electronic health record (EHR). The NIH Stroke Scale (NIHSS) at admission and discharge, and the modified Rankin Scale (mRS) at discharge, were assessed and documented by the stroke clinical team. Pre and post-procedural Alberta Stroke Program Early CT Score (ASPECT) scores were graded from neuroimaging by Dr. Aboul-Nour (board-certified vascular neurologist). Reperfusion status was determined at the time of thrombectomy by the neurointerventional attending using the Thrombolysis in Cerebral Infarction (TICI) score: TICI 0: no perfusion; TICI 1: penetration with minimal perfusion; TICI 2A: partial filling of < two-thirds of the vascular territory; TICI 2B: full territory filling with delayed flow; and TICI 3: complete perfusion. Only TICI 2B or higher patients were included.

### MicroVue Complement Analysis for Human Plasma

Complement testing was performed on human plasma samples using the MicroVue Complement Eight-Plex Panel (A901; Quidel) according to the manufacturer’s instructions. Complement proteins were captured and quantified on the surface of a 96- well microplate coated with antibodies specific for Bb, C3a, C4a, Factor I, Factor H, C5a, and sC5b9. Samples were diluted 1/100, and incubated for 2 hours. After washing unbound proteins, a biotinylated detection antibody cocktail was added and incubated for 1 hour. Next, streptavidin horseradish peroxidase was added and incubated for 20 minutes. The plate was developed with a substrate that reacts with HRP to generate a signal that is proportional to the amount of complement protein bound in the initial step. The plate was then measured on the Q-View Imager Pro (Quansys Biosciences). Data analyses were performed with the Q-View software ( Version 3.122). The software averages the duplicates, makes the necessary blank corrections, and calculates the best-fit non-linear regression curve for each standard. The concentration of the complement proteins in each samples was calculated from the standard curve. The final test result is dilution factor corrected.

### C3L ELISA for Human Plasma

Complement C3 level was measured with the Hycult C3 ELISA (HK366) per the manufacturer’s recommendation. Samples were diluted 1/16000 and incubated for 1 hour. After washing unbound proteins, a biotinylated detection antibody was added and incubated for 1 hour. Next, streptavidin horseradish peroxidase was added and incubated for 1 hour. The plate was developed with a TMB substrate that reacts with HRP to generate a signal that is proportional to the amount of complement protein bound in the initial step. The plate was then measured at 450nm with a plate reader. Data analyses were performed with SkanIt software (Version 5.0). The software averages the duplicates, makes the necessary blank corrections, and calculates the best-fit linear regression curve fit from the logarithmic transformed standards. The concentration of C3 in each sample was calculated from the standard curve. The final test result is dilution factor corrected.

### Statistical Analysis

All data are presented as mean ± standard deviation (SD) unless otherwise specified. Statistical analyses were performed using GraphPad Prism (version 10.2.2). All assumptions of normality and equal variance were tested prior to applying parametric tests. For comparisons between two groups, unpaired two-tailed Student’s t-tests were used. One-way ANOVA followed by Tukey’s post hoc test was applied for multi-group comparisons. Two-way ANOVA with Sidak post hoc tests was used for experiments involving two independent variables, such as treatment and time. Survival curves were analyzed using the log-rank (Mantel-Cox) test. For correlation analyses (e.g., IgG, IgM, and C3d relationships), Pearson correlation coefficients were calculated. Non-parametric tests (e.g., Mann–Whitney U test) were applied when the assumptions of normality were not met. For categorical data, including survival rates at day 5 between WT and C3 KO mice, Fisher’s exact test was used. p < 0.05 was considered statistically significant. The specific statistical tests used are indicated in the figure legends.

### Elastic Net Analysis

#### Data Preprocessing

Prior to analysis, all clinical variables were examined for distributional properties and completeness. Continuous variables were assessed for normality using Shapiro-Wilk tests and visually inspected. Non-normally distributed variables underwent appropriate transformations (log or square root) to improve normality. Missing data were not imputed; rather, observations with missing values were excluded from the analysis to avoid introducing potential biases associated with imputation methods. For categorical predictors, we set a minimum representation threshold of 10% in each level to ensure stable model estimation, that is the lowest ratio of each category must be > 1/10. Categories with insufficient representation were either recategorized (when conceptually appropriate) or excluded from further analyses to avoid model instability that can arise from sparsely populated levels and to ensure that each group contained a sufficient number of observations for robust estimation.

#### Statistical Modeling Strategy

We used elastic net regularization to investigate associations between acute glucose measurements, immune complement markers (e.g., C5a, C3a), and stroke outcomes (mRS_discharge and Post_ASPECTS). This method combines L1 (lasso) and L2 (ridge) penalties to address multicollinearity while selecting variables. The elastic net minimizes:

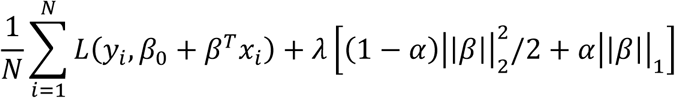

where L represents the negative log-likelihood function, λ controls overall penalty strength, and α determines the balance between ridge (α=0) and lasso (α=1) penalties. We examined two primary binary outcomes: Post-ASPECTS (defined as good when Post-ASPECTS > 5) and modified Rankin Scale at discharge (mRS_discharge, defined as good when ≤ 3). For each outcome, we explored a grid of candidate mixing parameters (α values) through 5-fold cross-validation. For binary outcomes, coefficients were converted into odds ratios and presented alongside the percent change in odds, for example, a negative percent change in odds (-X%), was interpreted as a decrease in the percent of “Good” outcome associated with a one-unit increase in that predictor. Model performance metrics including accuracy, sensitivity, specificity, positive predictive value (PPV), and negative predictive value (NPV) were calculated to assess each model’s discriminative ability. We assessed model performance using the area under the receiver operating characteristic curve (AUC) for binary outcomes and coefficient of determination (R²) for continuous outcomes.

We constructed models with increasing complexity. The baseline model incorporated clinical and demographic covariates to determine glucose’s importance based on predictive performance without complement factors. We then sequentially added individual complement factors to assess their independent contributions beyond baseline predictors. This approach allowed us to evaluate each complement marker’s incremental value while controlling for established clinical risk factors.

## Non-standard Abbreviations and Acronyms

A1c: Hemoglobin A1c
A_fib: Atrial fibrillation
ASPECT: Alberta Stroke Program Early CT Score
BBB: Blood–brain barrier
BMI: Body mass index
C3: Complement component 3
C3 KO: C3 knockout
DM: Diabetes mellitus
EB: Evans Blue
HG: Hyperglycemia
HLD: Hyperlipidemia
HT: Hemorrhagic transformation
HTN: Hypertension
IB4: Isolectin B4
L_MCA: Left middle cerebral artery
MCAO: Middle cerebral artery occlusion
MCV: Metabolic–Complement–Vascular
mRS: Modified Rankin Scale
NIHSS: National Institutes of Health Stroke Scale
PMI: Postmortem interval
R_MCA: Right middle cerebral artery
TICI score: Thrombolysis in Cerebral Infarction score
TNK: Tenecteplase
TTC: 2,3,5-Triphenyltetrazolium chloride
tPA: Tissue plasminogen activator
UEA-1: Ulex europaeus agglutinin I
WT: Wild Type

## Data availability

The clinical data used in this study are not publicly available due to patient privacy considerations but may be made available by the corresponding author upon reasonable request and with appropriate institutional approvals. Preclinical datasets supporting the findings of this study are also available from the corresponding author upon reasonable request.

## Materials availability

CR2-Crry was provided by Dr. Stephen Tomlinson and is available from his laboratory upon reasonable request, subject to a material transfer agreement.

## Code Availability

The custom code used to perform the open field behavior analysis is available at https://github.com/MA-121/openfield-behavior-analysis under the MIT License.

## Author Contribution

H.C. conceived the study and led the project execution, proposed the central hypothesis and collaborations, designed and performed the majority of experiments, conducted data analysis and interpretation, and wrote the manuscript. J.F. coordinated acquisition and analysis of human plasma samples and provided clinical input. C.T. arranged and processed human brain tissue, performed immunostaining and imaging. A.L. conducted predictive modeling and statistical analyses of human plasma datasets, including data interpretation. R.K. analyzed behavioral data and provided input for the manuscript. T.C. and A.K. assisted with behavioral experiments, blinding, and data collection. M.G. performed complement assays on human plasma samples. J.F.F., D.D., H.A-N., and N.M. collected clinical samples and associated patient data. S.T. provided the CR2-Crry reagent and contributed to study interpretation. L.D.M. contributed human brain samples and provided feedback and data interpretation. K.P. contributed human plasma samples and input. M.C., T.M.B., and G.K.S. jointly supervised the study, provided intellectual input and strategic oversight, overall research direction, data interpretation and contributed to writing the manuscript.

## Acknowledgments, Sources of Funding, & Disclosures

Acknowledgments:

We would like to thank Drs. Pardes Habib, Smarajit Mondal, Lindsey Druschel, for critical reading of the manuscript and feedback, and Christine Plant for help with manuscript submission.

## Supplemental Material

**Figure S1.**
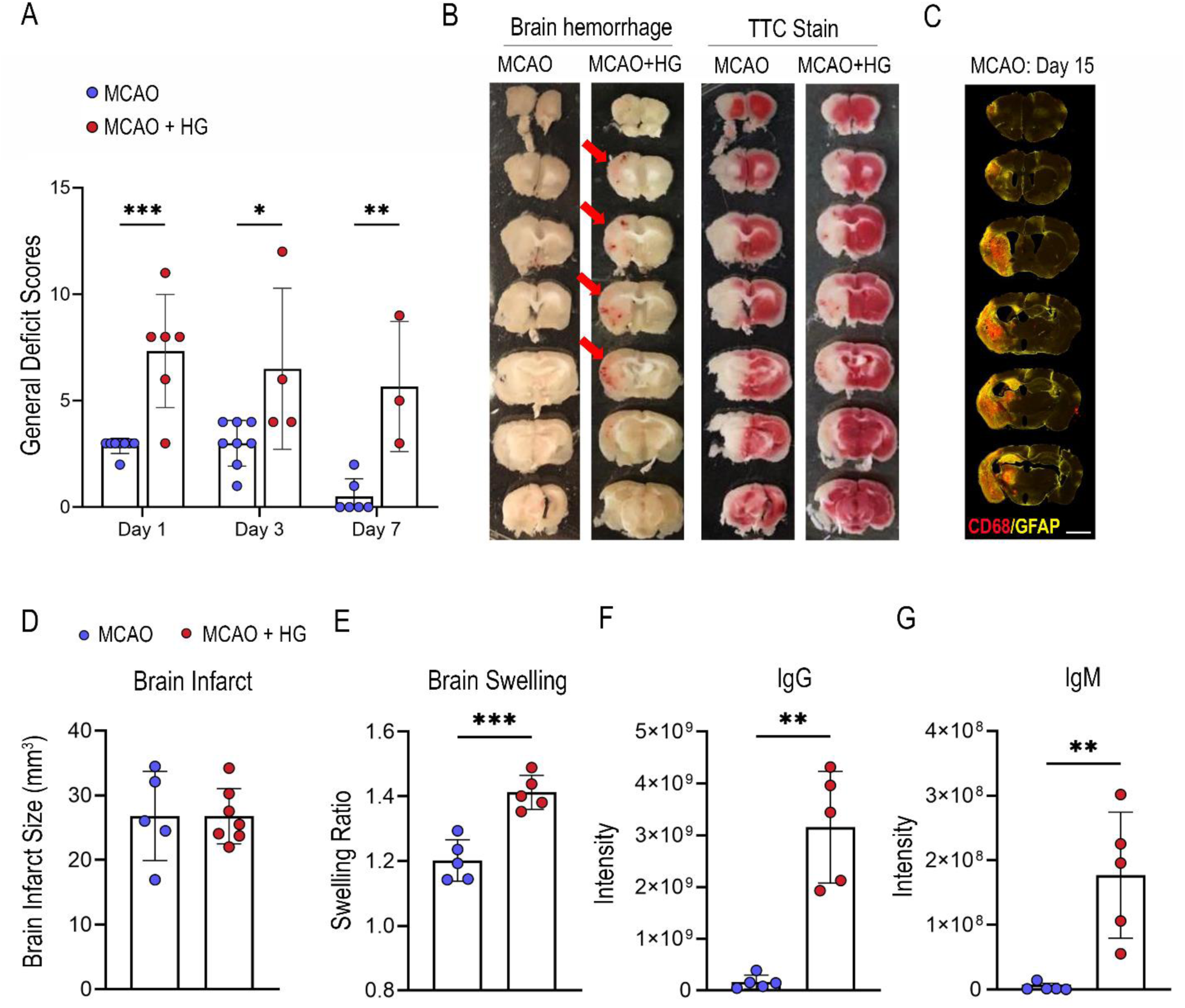
**A**, General neurological deficit scores in mice subjected to MCAO with or without acute hyperglycemia (HG). Mice were monitored after stroke, and scores were recorded in a blinded manner on days 1, 3, and 7 (n=6-8, two-way ANOVA). **B**, Representative post-mortem brain images from the same cohort as (A), showing spontaneous hemorrhagic transformation in hyperglycemic mice but not normoglycemic mice during the 14-day period; Representative TTC staining of the same brains confirmed the presence of infarcts, validating the successful stroke model. **C**, Representative immunostaining of CD68 and GFAP confirming successful induction of brain infarct in normoglycemic mice that survived. Scale bar: 2000 µm. **D**, Quantification of infarct volumes (from Fig.1K), n=5-7, unpaired t-test. **E**, Quantification of brain swelling at 24 hours. **F**, Quantification of IgG signal (from Fig. 1I). **G**, Quantification of IgM signal (from Fig. 1J). Data are shown as mean ± SD. *p<0.05, **p < 0.01, ***p < 0.001.

**Figure S2.**
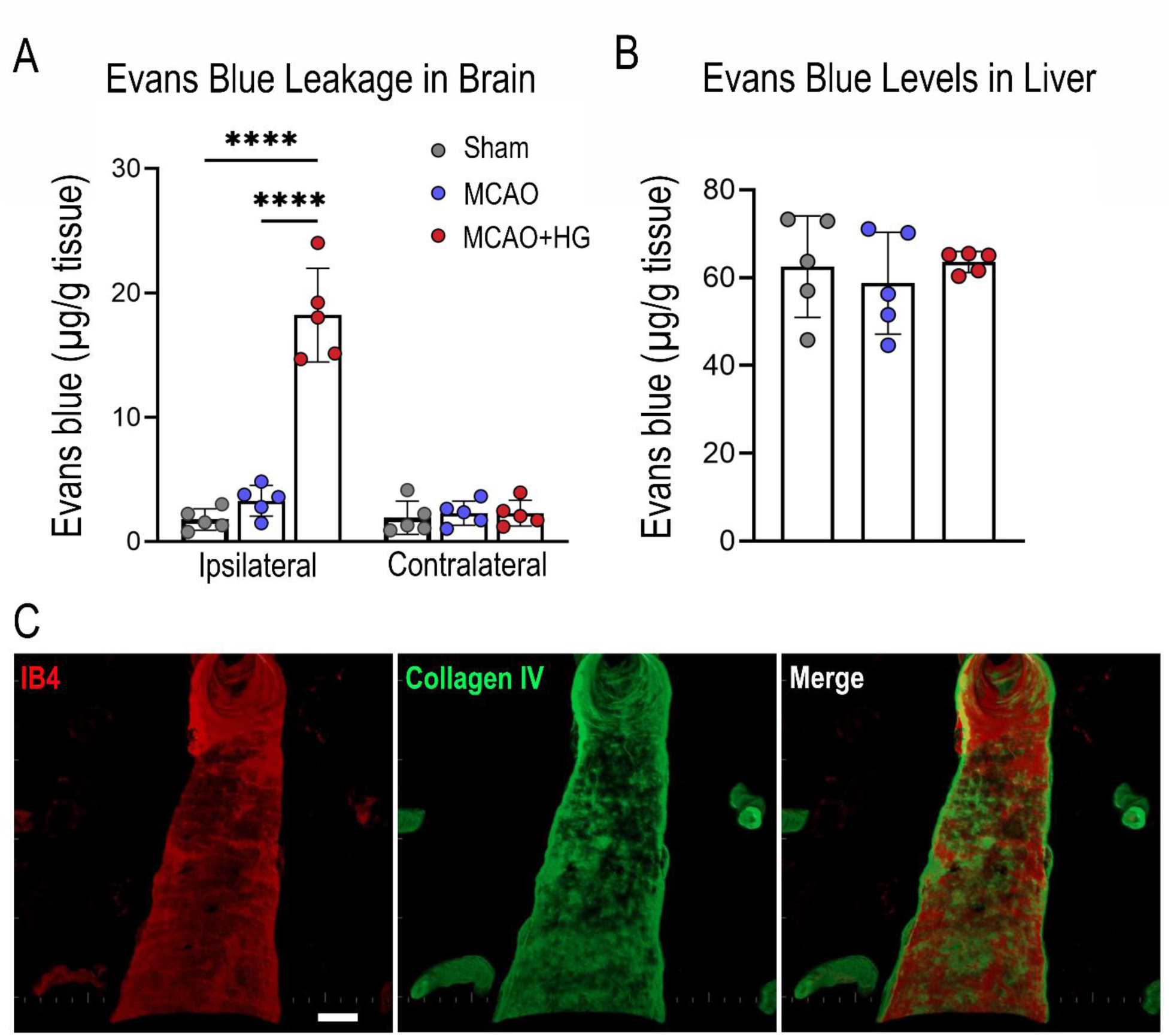
**A**, Quantification of Evans blue levels in brain tissues at 4.5 hours post- stroke (n = 5/group, ****p<0.0001). **B**, Quantification of Evans blue levels in liver tissue across Sham, MCAO, and MCAO + HG groups, showing no significant differences, confirming consistent systemic delivery of the tracer (n= 5/group). **C**, Representative longitudinal section showing co-staining of IB4 and collagen IV, demonstrating the predominantly luminal localization of IB4 staining in brain vessels (n = 5). Scale bar: 10µm.

**Figure S3.**
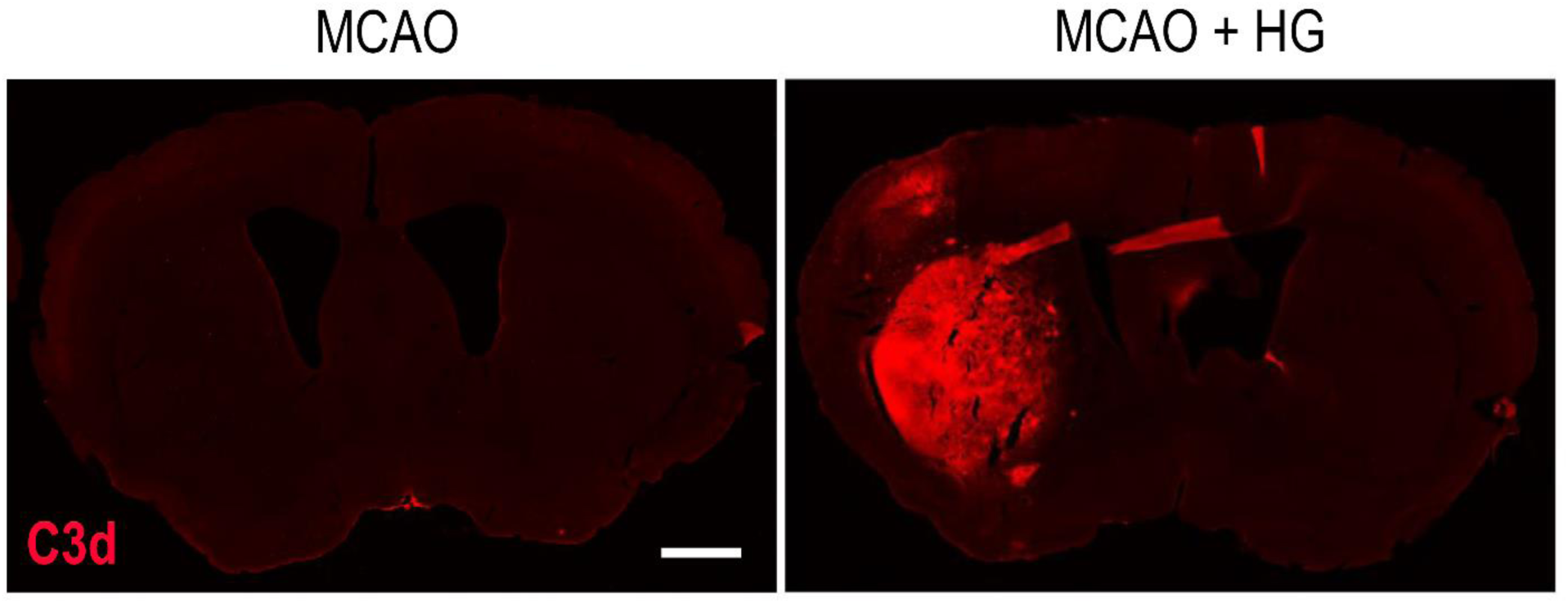
Representative brain sections from MCAO and MCAO + HG mice at 4.5 hours post-stroke stained for C3d (red). n=5-7. Scale bar:1000μm.

**Figure S4.**
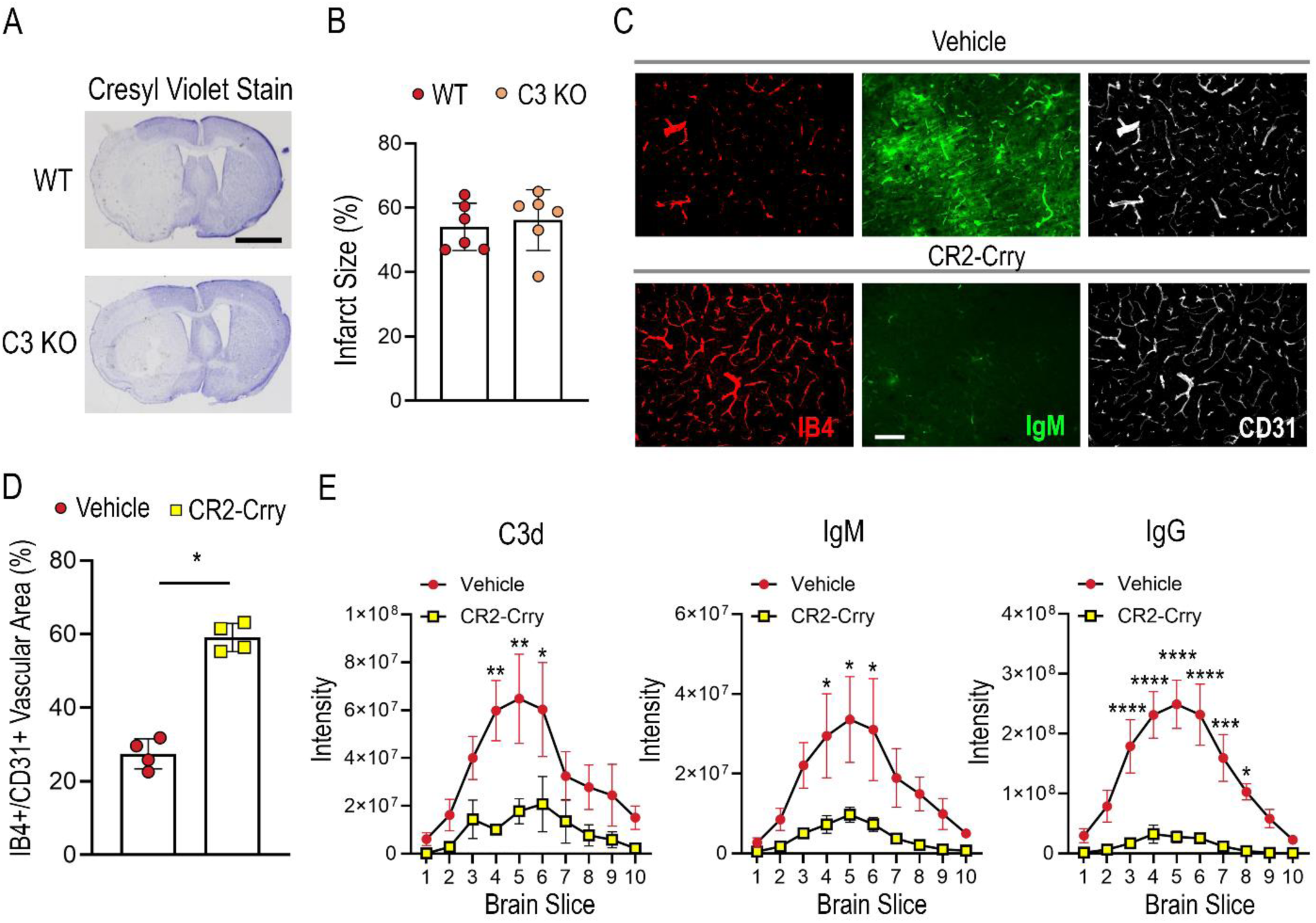
**A,** Cresyl violet staining of coronal brain sections from WT and C3 KO mice at 24 hours post-stroke. n=6/group. Scale bar: 2000µm. **B,** Quantification of infarct size revealed no significant difference between groups. n=6/group. Data are presented as mean±SD. **C-D**, CR2-Crry treatment increased IB4⁺/CD31⁺ vascular coverage compared to vehicle-treated mice. Scale bar: 100µm. n=4/group. Unpaired t-test, *p<0.05. Data are presented as mean±SD. **E**, Distribution plots showing brain slice-wise quantification of C3d, IgM, and IgG intensities in vehicle vs. CR2–Crry–treated mice. n=4/group. *p<0.05; **p<0.01; ***p<0.001, ****p<0.0001. Two-away ANOVA followed by Šídák’s multiple comparisons test. Data are presented as mean± SEM.

**Figure S5.**
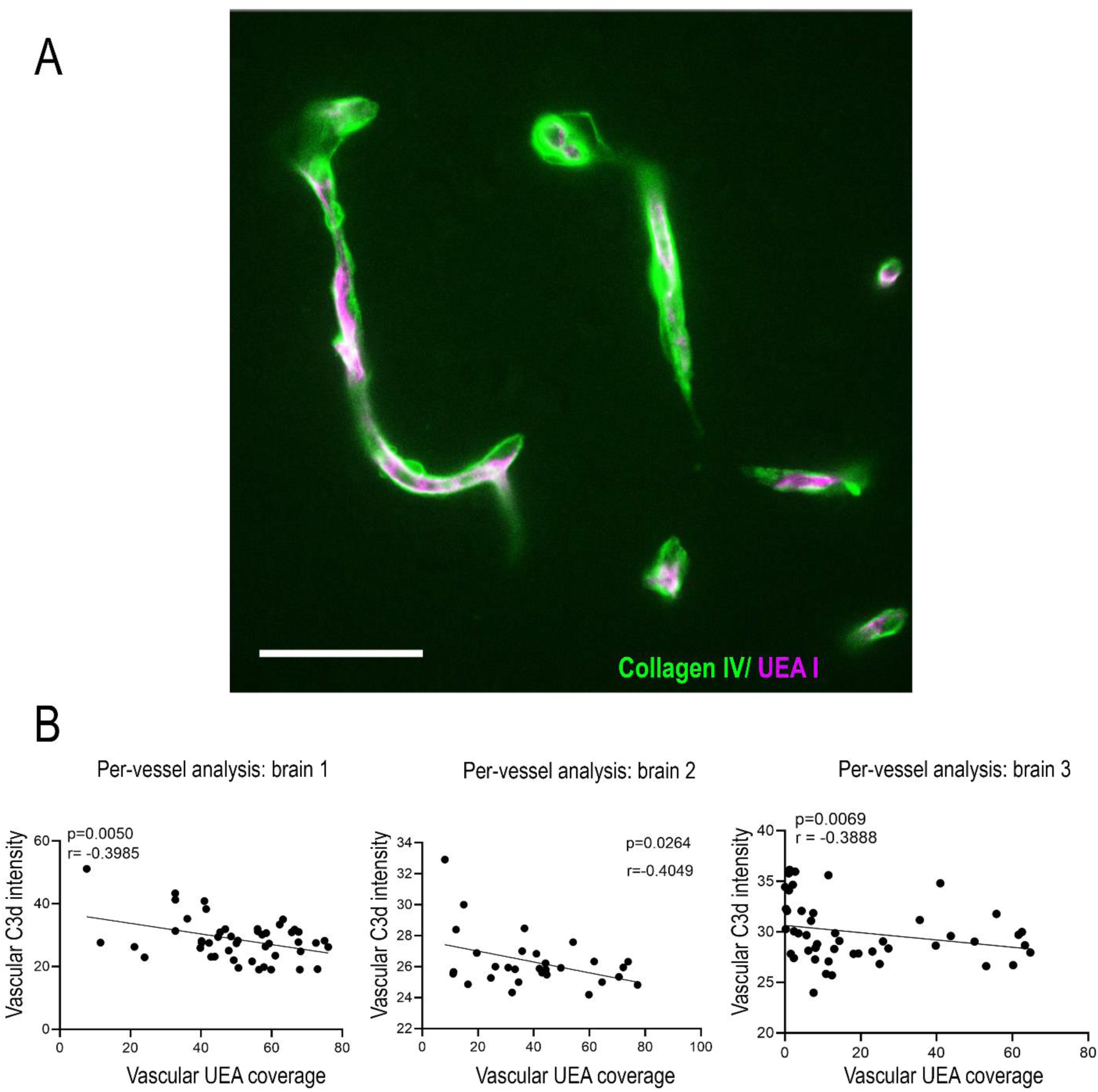
**A**, Representative confocal image showing luminal localization of UEA I staining in human brain vessels (Collagen IV+). Scale bar: 50µm. **B**, Per-vessel correlation analysis of vascular C3d intensity and UEA I coverage in three independent hyperglycemic stroke brains. Each dot represents an individual vessel. Correlations were assessed using Pearson or Spearman methods, depending on data distribution normality.

**Figure S6.**
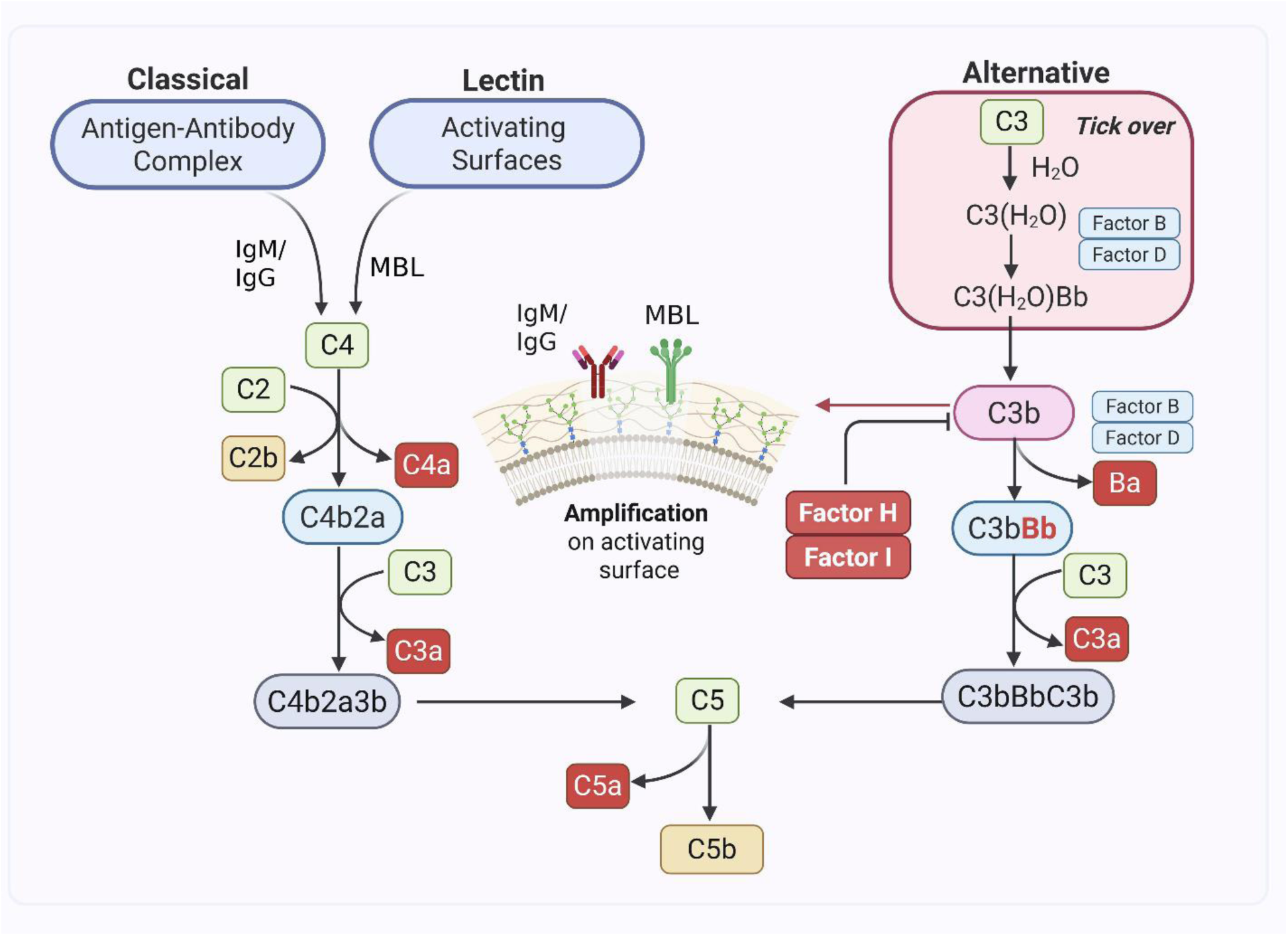
Complement activation pathways and their relationship to vascular surface damage. Red-colored components are the soluble complement products detected in plasma in this study cohort.

**Figure S7.**
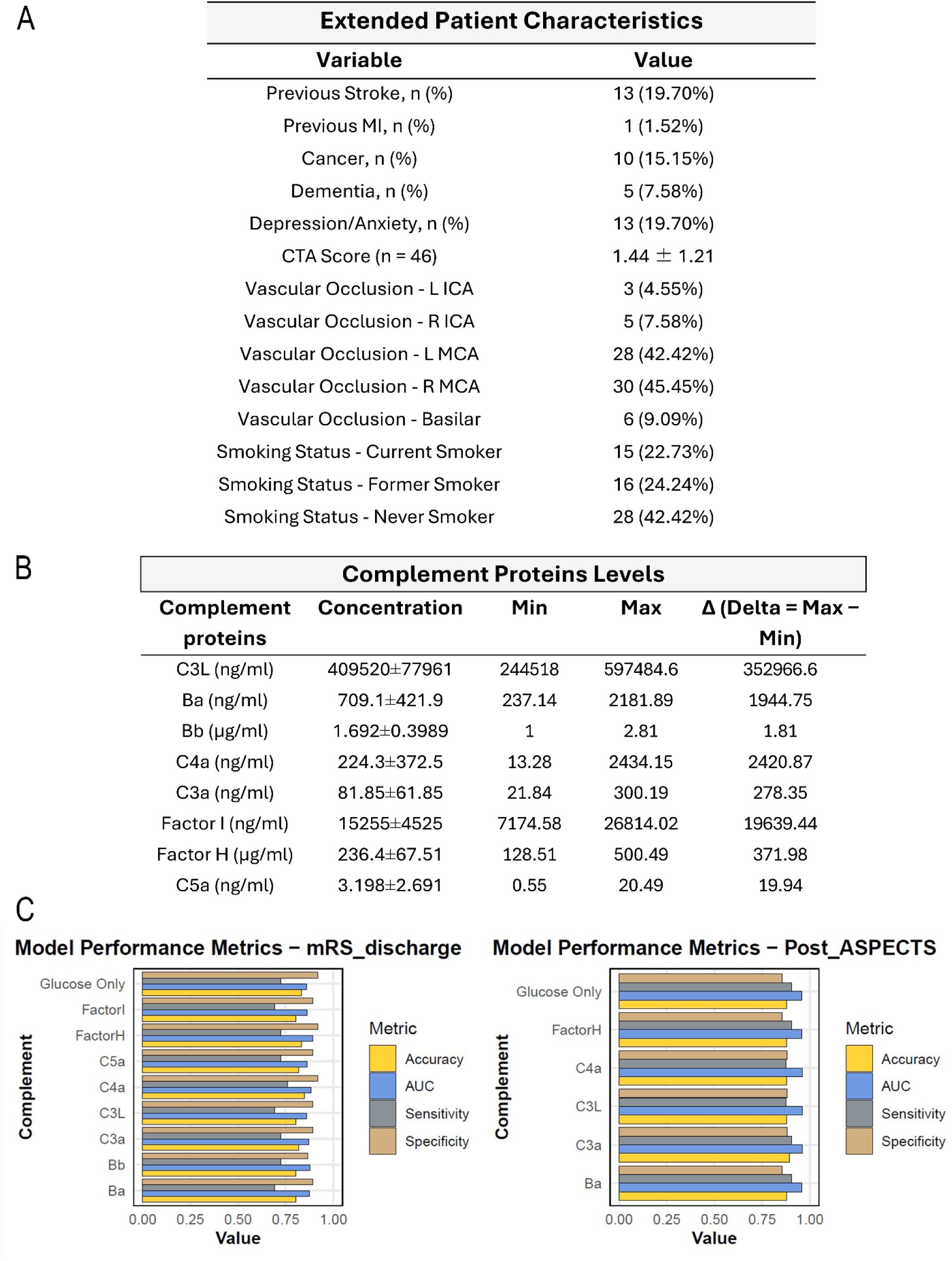
**A**, Extended demographic and clinical characteristics of stroke patients included in the plasma cohort. **B**, Distribution ranges of plasma complement activation products measured in the cohort. **C**, Performance of the Elastic Net regression model in predicting post-thrombectomy outcomes based on complement and clinical variables.

**Table S1. Demographic and Clinical Characteristics of the Post-mortem Brain Cohort. Abbreviations: PMI, postmortem interval; DM, diabetes mellitus; A1c, hemoglobin A1c; HTN, hypertension; BMI, body mass index.**

**Table S2. Clinical Predictors of Outcome Identified by Elastic Net Regression. Abbreviations: HTN, hypertension; HLD, hyperlipidemia; DMII, type 2 diabetes mellitus; A_fib, atrial fibrillation; A1c, hemoglobin A1c; tPA_TNK, tissue plasminogen activator or tenecteplase; L_MCA, left middle cerebral artery; R_MCA, right middle cerebral artery.**

